# Derlin-mediated ERAD of lipid regulator ORMDL3 safeguards mitochondrial function

**DOI:** 10.64898/2026.02.27.708653

**Authors:** Nicola A. Scott, Jeremiah M. Afolabi, Andrea G. Marshall, Jenny C. Schafer, Victoria R. Baskerville, Praveena Prasad, Ashlesha A. Kadam, Catherine Som de Cerff, Thomas Whisenant, Mark A. Phillips, Dhanendra Tomar, Melanie R. McReynolds, Antentor Hinton, Sonya E. Neal

**Affiliations:** School of Biological Sciences, Department of Cell and Developmental Biology, University of California San Diego, La Jolla, CA 92093, USA; Department of Molecular Physiology and Biophysics, Vanderbilt University, Nashville, TN, 37232, USA; Department of Cell and Developmental Biology, Vanderbilt University, Nashville, TN, 37232, USA; Department of Medicine, Division of Clinical Pharmacology, Vanderbilt University Medical Center, Nashville, TN, 37232, USA; Department of Biochemistry and Molecular Biology, The Huck Institute of the Life Sciences, Pennsylvania State University, State College, PA 16802, USA; Department of Internal Medicine, Wake Forest University School of Medicine, Winston-Salem, NC 27109, USA; Center for Computational Biology and Bioinformatics, University of California San Diego, La Jolla, CA 92093, USA; Department of Integrative Biology, Oregon State University, Corvallis, OR 97331, USA; Howard Hughes Medical Institute, Chevy Chase, MD 20815, USA

**Keywords:** Derlins, ERAD, MERCs, ER, mitochondria, ORMDLs

## Abstract

Mammalian Derlin proteins (Derlin-1, Derlin-2, and Derlin-3) are conserved components of the endoplasmic reticulum–associated degradation (ERAD) machinery that mediate the retrotranslocation and proteasomal degradation of misfolded ER-resident proteins. However, their paralog-specific contributions to cellular homeostasis remain poorly understood. Here, we show that Derlin deficiency disrupts mitochondrial architecture and results in mitochondrial fragmentation and tightening of ER-mitochondria contact sites (MERCs) in HEK293 cells. Mechanistically, we identify ORMDL proteins, evolutionarily conserved negative regulators of sphingolipid biosynthesis, as substrates of Derlin-2- and Derlin-3-dependent ERAD. Derlin deficiency leads to selective accumulation of ORMDL3 and its dose-dependent enrichment at MERCs, where it drives mitochondrial dysfunction in respiration and calcium handling. Reducing ORMDL3 levels restores mitochondrial function, establishing ORMDL3 as a key effector downstream of Derlin loss. Our work establishes ERAD as a critical mechanism of protein quantity control that safeguards organelle homeostasis by preventing aberrant accumulation and mislocalization of ER clients at inter-organelle contact sites. Given that ORMDL family members are central regulators of sphingolipid metabolism and are genetically linked to inflammation, cancer, asthma, inflammatory bowel disease, type 1 and type 2 diabetes, multiple sclerosis, obesity, and nonalcoholic fatty liver disease, these findings connect ERAD-dependent spatial control to sphingolipid homeostasis and a broad spectrum of human pathologies.

## Introduction

The endoplasmic reticulum (ER) houses an elaborate protein quality control network that ensures high-fidelity protein synthesis and folding (Adams et al., 2019). Proteins that fail to attain their native conformation are retained within the ER and eliminated to prevent proteotoxic stress. A principal mechanism for this clearance is ER-associated degradation (ERAD), an evolutionarily conserved pathway in which misfolded proteins are retrotranslocated from the ER to the cytosol and degraded by the proteasome (Needham & Brodsky, 2013).

All ERAD pathways rely on substrate recognition and ubiquitination by dedicated E3 ubiquitin ligases and require the AAA+ ATPase Cdc48 (p97 in mammals) to extract misfolded proteins from the ER membrane or to drive the retrotranslocation of substrates across the membrane for proteasomal degradation (Bays et al., 2001; Bodnar & Rapoport, 2017; Neal et al., 2017; Twomey et al., n.d.; Ye et al., 2005). Central to this process are Derlins, ER-resident, multi-pass membrane proteins classified as rhomboid pseudoproteases and core components of the ER protein quality control machinery (Greenblatt et al., 2011). In yeast, two Derlins, Der1 and Dfm1, mediate the retrotranslocation of luminal and membrane ERAD substrates, respectively (Knop et al., 1996; Neal et al., 2018). Notably, mammals encode three paralogs, Derlin-1, Derlin-2, and Derlin-3, all of which have been implicated in ERAD (Kadowaki et al., 2018; Lilley & Ploegh, 2004a; Sato & Hampton, 2006). Human Derlin-1, the closest human homolog of Dfm1, was initially identified through its role in cytomegalovirus-mediated degradation of MHC I (Lilley & Ploegh, 2004b). Subsequent biochemical studies have identified numerous Derlin substrates, including CFTR and CFTR-ΔF508, ENaC, proinsulin, sonic hedgehog, α1-antitrypsin (NHK mutant), and viral nonstructural proteins, underscoring the broad physiological relevance of Derlin-dependent ERAD (Hoelen et al., 2015; Huang et al., 2013; Oda et al., 2006; You et al., 2017).

Despite their apparent functional overlap, accumulating evidence suggests that Derlin paralogs also perform non-redundant roles *in vivo*. Global knockout of Derlin-1 or Derlin-2 in mice results in embryonic or perinatal lethality, whereas Derlin-3 knockout mice develop normally, consistent with its restricted tissue expression (Dougan et al., 2011; Eura et al., 2012). Conditional knockout studies further demonstrate that Derlin-1 and Derlin-2 are essential in highly secretory or proteotoxically stressed tissues, including the kidney and neurons (Ren et al., 2018; Sugiyama et al., 2021). Notably, Derlin-2 loss uniquely causes severe ER stress and pathology in podocytes, skeletal dysplasia due to defective collagen secretion in chondrocytes, and peripheral neuropathy in Schwann cells (Volpi et al., 2019). In cancer, Derlin paralogs also display distinct expression patterns: Derlin-1 is elevated in breast and colon cancers (Tan et al., 2015; J. Wang et al., 2008), whereas Derlin-3 is enriched in lung and kidney cancers (Lin et al., 2025; Niu et al., 2025), suggesting paralog-specific roles in stress adaptation and disease.

Despite these insights, a systematic cellular comparison of Derlin paralogs has been lacking. Here, we performed a comprehensive characterization of Derlin-1, Derlin-2, and Derlin-3 using CRISPR-engineered knockout HEK293 cells. We find that loss of Derlin-2, but not Derlin-1 or Derlin-3, induces pronounced ER stress, identifying Derlin-2 as the dominant paralog in maintaining ER proteostasis. Unexpectedly, loss of Derlin-1 predominantly perturbs ER morphology, whereas loss of all three Derlin paralogs results in pronounced mitochondrial fragmentation. Derlin-2 KO cells, and to a lesser extent Derlin-3 KOs, exhibit marked mitochondrial dysfunction, including impaired calcium handling, reduced respiratory capacity, and disrupted mitochondria–ER contact sites (MERCs). These organelle-specific defects are accompanied by distinct lipidomic, proteomic, and metabolomic remodeling across the Derlin knockout backgrounds.

Mechanistically, we identify a paralog-specific role for Derlins in spatially limiting the ERAD substrate ORMDL3, a negative regulator of sphingolipid biosynthesis, from aberrant accumulation at MERCs. In the absence of Derlin-2, ORMDL3 escapes ERAD, redistributes from the ER to MERCs, and disrupts mitochondrial respiration and calcium flux. Although ORMDL1 and ORMDL2 are also ERAD substrates, their stabilization does not result in MERC mislocalization, revealing functional divergence among ORMDL paralogs at the cellular level. Together, our findings uncover distinct roles of each Derlin paralog in mediating cellular homeostasis and provide mechanistic insight into why ORMDL3, but not other ORMDL family members, is disproportionately implicated in diverse diseases, including cancer, inflammation, diabetes, and asthma (Brown & Spiegel, 2023; Davis et al., 2018a; Gupta et al., 2015; H. Wang et al., 2025).

## Results

Derlin-1, Derlin-2, and Derlin-3 are ER-resident proteins predicted to contain six transmembrane domains and share structural similarity within the rhomboid-like superfamily (**Fig. 1A**) (Bhaduri, Scott, et al., 2022). Despite this conservation, Derlin-1 and Derlin-2 possess a cytosolic SHP motif in their C-terminal tail that mediates recruitment of the AAA-ATPase p97, a central component that powers the ERAD process via ATP hydrolysis, whereas Derlin-3 lacks lack this motif (Greenblatt et al., 2011; Hoelen et al., 2015). To establish whether the Derlin paralogs exhibit distinct foundational properties, we first undertook a comprehensive characterization of each. We tested whether the three mammalian Derlins are required for alleviating ER stress, which is known to activate the unfolded protein response (UPR). We utilized previously generated HEK293 cell lines in which each Derlin paralog was individually knocked out (Derlin-1 KO: D1 KO, Derlin-2 KO: D2 KO, and Derlin-3 KO: D3 KO), as well as a triple-knockout line lacking all three Derlins (Derlin TKO: D TKO) generated using CRISPR–Cas9. Successful depletion of Derlin-1 and Derlin-2 paralog was confirmed by immunoblotting (**Fig. S1A**). Due to the lack of reliable antibodies against endogenous Derlin-3, we employed a TOPO cloning strategy to amplify and sequence the Derlin-3 locus in Derlin-3 KO and Derlin TKO cells and confirmed the presence of a Derlin-3 allele that was consistent with previously published work that generated these lines (**Fig. S1B**) (Kadowaki et al., 2015). Strikingly, loss of Derlin-2 led to robust induction of multiple UPR components, including BiP, PERK, IRE1α, and ATF6. In contrast, Derlin-1 KO and Derlin-3 KO cells exhibited little to no comparable upregulation, whereas Derlin TKO cells showed the highest induction of UPR components, consistent with partial redundancy among Derlin family members (**Fig. 1B**). Ectopic expression of Derlin-2 in the knockout background restored ER stress marker levels to those observed in WT cells (**Fig. 1C**), demonstrating that Derlin-2 is the major Derlin paralog required for alleviating ER stress.

**Figure 1.**
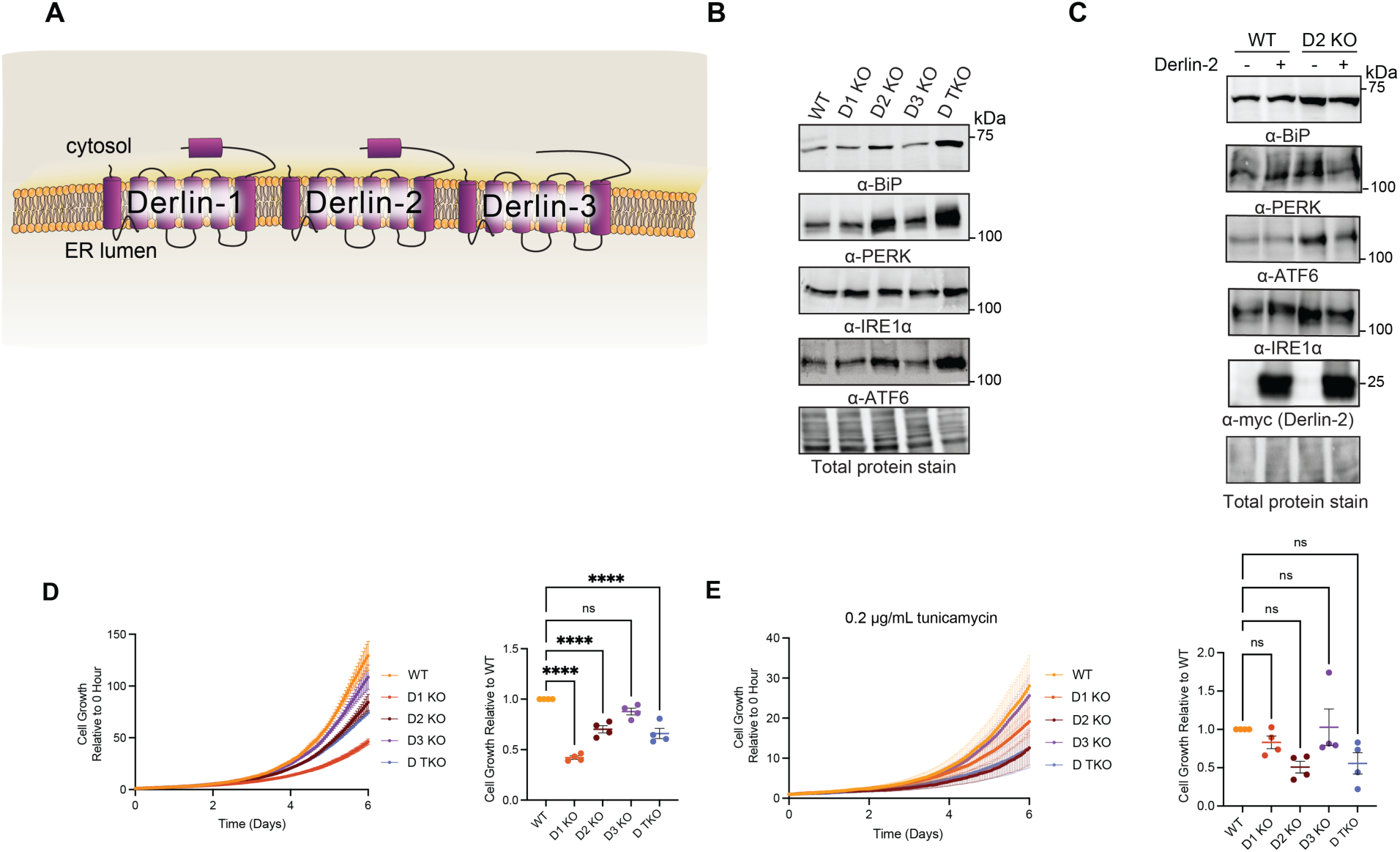
Loss of each Derlin confers distinct sensitivity to ER stress and differentially affects cell proliferation. **(A)** Schematic of mammalian Derlin proteins. Derlin-1, Derlin-2, and Derlin-3 are ER membrane proteins with six predicted transmembrane domains. Derlin-1 and Derlin-2 contain a C-terminal SHP motif that recruits p97, whereas Derlin-3 lacks this feature, but can still recruit p97. **(B)** Immunoblot analysis of ER stress markers (BiP, PERK, IRE1α, ATF6) in HEK293 wild-type (WT), Derlin-1 KO, Derlin-2 KO, Derlin-3 KO, and Derlin triple KO cells. Representative of *n* = 3 independent experiments. **(C)** Rescue of ER stress by ectopic expression of *DERLIN2.* HEK293 WT and Derlin-2 KO cells were transfected with Derlin-2–Myc/His for 24 h, followed by immunoblotting for BiP, PERK, IRE1α, and ATF6. Representative of n = 4 independent experiments. **(D)** Proliferation curves of HEK293 WT and *DERLIN* KO cell lines measured by live-cell confluence every hour for 6 days. Confluence was normalized to each cell line’s 0 h value. Area under the curve was quantified and compared to WT. Representative of n = 4 independent experiments. Statistical significance was determined by one-way ANOVA followed by Tukey’s multiple-comparison test. ns, not significant; ****P < 0.0001. **(E)** Proliferation of HEK293 WT and Derlin KO cells treated with 200 ng/mL tunicamycin. Confluence was measured as in (D). Area under the curve for each cell line was quantified and compared to WT. Representative of *n* = 4 independent experiments. Statistical significance was determined by one-way ANOVA followed by Tukey’s multiple-comparison test. ns, not significant.

We next investigated whether Derlin deficiency impacts cell proliferation. Real-time confluence monitoring over multiple days revealed that Derlin-1 KO, Derlin-2 KO, and Derlin TKO cells proliferated significantly more slowly than WT, whereas growth of Derlin-3 KO cells was comparable to WT (**Fig. 1D**). Quantification of the growth curves confirmed reduced proliferative capacity in Derlin-1 KO, Derlin-2 KO, and Derlin TKO cells, indicating that both Derlin-1 and Derlin-2 are required for optimal cell growth. Given the activation of ER stress in Derlin-2 KO and Derlin TKO cells, we hypothesized that these mutant lines would exhibit increased sensitivity to ER stress. Under vehicle treatment, cells displayed the expected growth profiles across genotypes (**Fig. S1C**). Upon tunicamycin treatment, WT including all single KOs and TKOs had decreased growth in comparison to their vehicle treatment. However, although not statistically significant, Derlin-2 KO and Derlin TKO cells trended towards the greatest reduction in growth, with Derlin-1 KO cells exhibiting an intermediate response relative to WT. In contrast, WT and Derlin-3 KO cells displayed similar growth under ER stress conditions. These results are consistent with increased ER stress sensitivity in Derlin-2–deficient cells (**Fig. 1E**). Together, these data reveal distinct effects for the Derlin paralogs, with Derlin-2 as the major contributor for alleviating ER stress, while both Derlin-1 and Derlin-2 are necessary to support cell proliferation.

### Derlin loss leads to fragmented mitochondria and alterations in mitochondria-ER contact sites

Given that each Derlin knockout exerts distinct effects on ER stress and cell proliferation, we next examined how Derlins influence cellular organization at a broader level, with particular emphasis on the ER, as disruptions in ER homeostasis are known to profoundly alter ER morphology. To address this, we employed transmission electron microscopy (TEM) to quantify ER area and perimeter (**Fig. 2A**, *highlighted in orange*). Notably, only Derlin-1 KO and Derlin TKO showed a significant increase in ER area and perimeter compared to WT (**Fig. S1D**). Unexpectantly, we also observed pronounced ultrastructural alterations in mitochondria, particularly in mitochondrial length and area (**Fig. 2A**, *highlighted in purple and red respectively*). Loss of any individual Derlins resulted in a striking reduction in mitochondrial length and area relative to WT cells (**Fig. 2B-D**). In addition, Derlin-1 KO and Derlin-3 KO cells exhibited significantly reduced mitochondrial area (**Fig. 2B, D**), whereas Derlin TKO cells showed a more pronounced decrease in both mitochondrial length and area compared with the single KO lines (**Fig. 2B-D**). As controls, we analyzed knockdowns of established regulators of mitochondrial dynamics, including fusion proteins (OPA1 and MFN2) and cristae-organizing components (CHCHD3, CHCHD6, and Mitofilin), which displayed the expected mitochondrial fragmentation phenotype (**Fig. S2A-D**) (An et al., 2012; Cipolat et al., 2004). To further support that loss of Derlins leads to mitochondrial fragmentation, three-dimensional surface renderings of mitochondria were reconstructed from live-cell imaging of mitochondrial dynamics using MitoTracker staining. Loss of each Derlin resulted in marked reductions in mitochondrial area and volume, accompanied by a concomitant decrease in mitochondrial sphericity, consistent with a fragmented mitochondrial morphology (**Fig. 2F-I**). We next quantified the distance between MERCs (**Fig. 2A**, *in yellow*), a critical hub for lipid biosynthesis and Ca²⁺ signaling that is known to function optimally within a defined spatial range (Dematteis et al., 2024). Consistent with previous reports, knockdown of the mitochondrial dynamics proteins MFN2 and OPA1 resulted in a marked increase and decrease in MERC distance, respectively, compared with WT cells. Quantitative analysis of MERC spacing in Derlin knockout cells revealed a significant reduction in contact distance in Derlin-2 KO, Derlin-3 KO, and Derlin TKO cells, whereas Derlin-1 KO cells exhibited an increase in MERC distance (**Fig. 2E**). Together, these findings demonstrate that loss of any Derlin paralog disrupts organelle ultrastructure and morphology. Specifically, Derlin-1 knockout cells exhibit altered ER morphology, all single Derlin knockouts display mitochondrial fragmentation, and Derlin-2 and Derlin-3 knockouts show reduced MERC distance.

**Figure 2.**
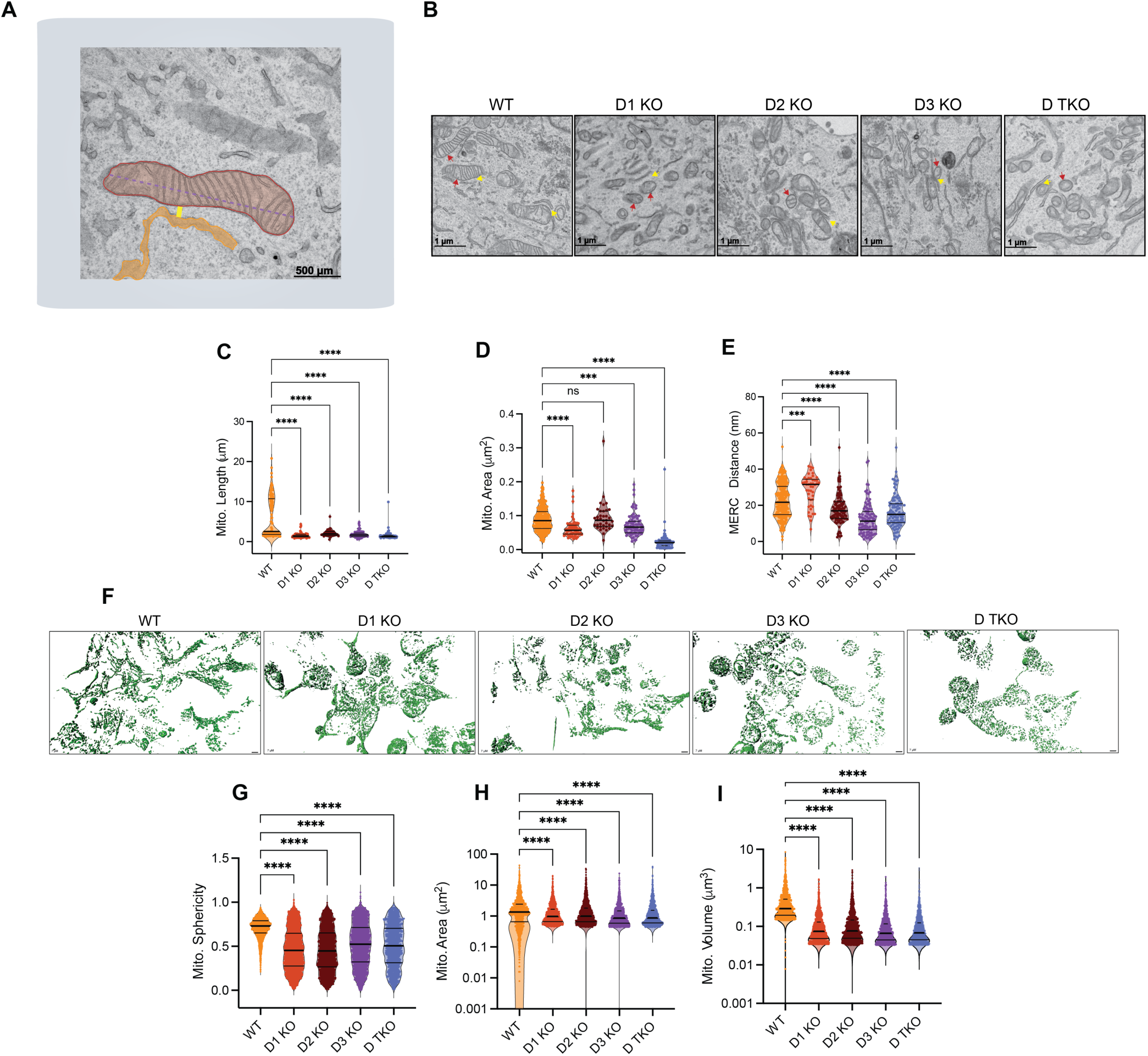
Transmission electron microscopy reveals mitochondrial remodeling following disruption of Derlins. **(A)** Representative TEM image illustrating measurement of mitochondria area (*red*), length (*purple*), ER area (*orange*), and MERCs distance (*yellow*). **(B)** Representative TEM images are shown for (A) WT, Derlin-1 KO, Derlin-2 KO, Derlin-3 KO, Derlin TKO. Mitochondria display distinct alterations in size and shape depending on the targeted pathway: red arrows indicate mitochondria; yellow arrows MERCs; scale bars, 2 µm. **(C-E)** Quantitative analyses of mitochondrial length (C), mitochondrial area (D), and mitochondria–ER contact (MERC) distance (E) were performed from TEM images using standardized criteria across all conditions. Each data point represents an individual mitochondrion. Statistical significance was determined using one-way ANOVA with multiple-comparison correction. ***P < 0.001, ****P < 0.0001; ns, not significant. **(F)** Representative 3D surface renderings of mitochondria reconstructed from high-resolution confocal z-stack imaging are shown for WT, Derlin-1 KO, Derlin 2 KO, Derlin 3 KO, and Derlin TKO in HEK293 cells. Mitochondria are displayed as green surface renderings following automated segmentation and batch surface generation, with identical parameters across all treatment groups; scale bar, 7 µm. **(G-I)** Quantification of mitochondrial sphericity (G), mitochondrial area (H), and mitochondrial volume **(I)**. Each dot represents an individual mitochondrion. Conpared with WT, all Derlin KOs produce pronounced reductions in mitochondrial area and volume and a shift toward increased sphericity, consistent with mitochondrial fragmentation. Statistical significance was determined using one-way ANOVA with appropriate multiple-comparison correction. ****P < 0.0001; ns, not significant.

We next asked whether ER stress itself could phenocopy these mitochondrial defects since our results above showed Derlin-2 KO induced ER stress and activated UPR. Surprisingly, tunicamycin and thapsigargin treatment induced the opposite effect, producing elongated, hyperfused mitochondrial networks rather than the fragmented morphology observed in Derlin-2 KO cells (**Fig. S2E-H**). These findings indicate that the mitochondrial defects observed in Derlin KO cells are not secondary to ER stress but instead represent a direct consequence of Derlin loss.

### Derlin-2 and Derlin-3 are required for mitochondrial function

Given the profound mitochondrial morphological defects observed in Derlin KO cells, we asked whether these changes impair mitochondrial function. Seahorse extracellular flux analysis revealed that Derlin-2 KO and Derlin-3 KO cells exhibited a marked reduction in maximal respiration and spare respiratory capacity, with Derlin TKO cells showing the most pronounced defects compared with WT (**Fig. 3A-B**). In contrast, Derlin-1 KO cells displayed minimal to no detectable changes in all respiratory parameters examined, despite exhibiting clear mitochondrial morphological abnormalities (**Fig. 3A-B**). Mitochondrial fragmentation and MERCs abnormalities are frequently associated with defects in Ca²⁺ shuttling between the ER and mitochondria(Dematteis et al., 2024). We therefore examined the impact of Derlin deficiency on ER and mitochondrial Ca²⁺ flux (**Fig. 3C**). In permeabilized HEK293 cells treated with thapsigargin to inhibit ER Ca²⁺ reuptake, ER Ca²⁺ release was unchanged across all Derlin KO lines, indicating that Derlin loss does not impair ER Ca²⁺ release (**Fig. S3A**). Given that MERCs tightening observed in Derlin-2 KO and Derlin-3 KO cells (**Fig. 2E**) could disrupt Ca²⁺ transporter organization and function, we next measured mitochondrial Ca²⁺ efflux and uptake. Following acute inhibition of mitochondrial Ca²⁺ uptake using the mitochondrial calcium uniporter inhibitor Ru360, Derlin-2 KO cells exhibited a significant reduction in the rate of mitochondrial Ca²⁺ efflux compared with WT, measured (**Fig. 3D**). Prior to Ru360 addition, Derlin-2 KO and Derlin-3 KO cells exhibited a trend towards reduced mitochondrial Ca²⁺ uptake that did not reach statistical significance; however, with Derlin TKO cells, mitochondrial Ca²⁺ uptake was significantly attenuated, which may be due to the combined loss of Derlin-2 and Derlin-3 (**Fig. 3E**).

**Figure 3.**
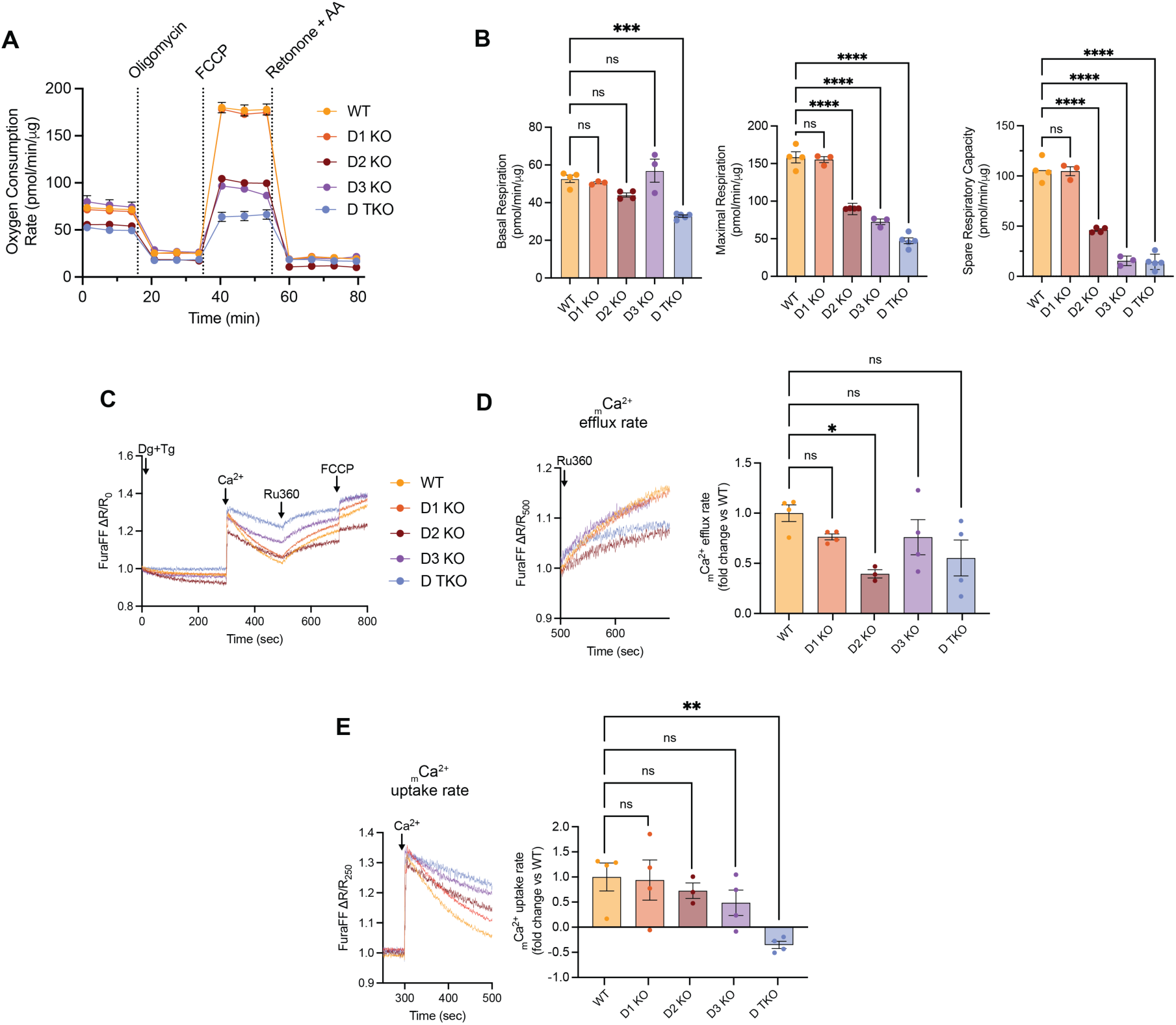
Derlins influence mitochondrial respiration and Ca²⁺ homeostasis. **(A)** Oxygen consumption rate (OCR) was measured in HEK293 WT, Derlin-1 KO, Derlin-2 KO, Derlin-3 KO, and Derlin TKO cells using a Seahorse Analyzer. **(B)** Quantification of basal respiration, maximal respiration, and spare respiratory capacity was quantified from (A). Data represent *n* = 3–4 independent experiments (mean ± SEM). Statistical significance was determined by one-way ANOVA followed by Tukey’s multiple-comparison test. ***P < 0.001, ****P < 0.0001; ns, not significant. **(C)** Ratiometric monitoring of mitochondrial Ca²⁺ dynamics in HEK293 WT and Derlin KO cells using FuraFF. **(D)** Mitochondrial Ca²⁺ (mCa²⁺) efflux rate was normalized at 500 sec. Fold change in mCa²⁺ efflux of Derlin KO cells was compared to WT. Data represent *n* = 3–4 independent experiments (mean ± SEM). Statistical significance was determined using one-way ANOVA followed by Dunnett’s multiple-comparison test. *P < 0.05; ns, not significant. **(E)** mCa²⁺ uptake rate was normalized at 300 sec. Fold change in mCa²⁺ uptake of Derlin KO cells was compared to WT. Data represent *n* = 3–4 independent experiments (mean ± SEM). Statistical significance was determined using one-way ANOVA followed by Dunnett’s multiple-comparison test. **P < 0.01; ns, not significant.

To further validate mitochondrial defects associated with Derlin loss, particularly in Derlin-2 and Derlin-3 KO cells, we performed global metabolomic profiling across all Derlin KO lines. Principal component analysis revealed that Derlin-2 KO and Derlin-3 KO cells clustered together and segregated distinctly from Derlin-1 KO and Derlin TKO cells (**Fig. 4A**). In addition, Derlin-1 KO cells exhibited the most extensive metabolic upregulation, with pronounced enrichment of metabolites associated with D-amino acid metabolism, and porphyrin metabolism relative to Derlin-2 and Derlin-3 KO cells (**Fig. S4A-B**). Hierarchical clustering of metabolite abundance revealed a marked reduction in methylnicotinamide in Derlin-2 KO cells (**Fig. 4B**), indicative of diminished NAD⁺ salvage pathway intermediates, as reduced levels of this intermediate have been previously associated with lowered NAD⁺ availability, mitochondrial fragmentation, and mitochondrial dysfunction (Su et al., 2024). Guided by this finding, we performed targeted quantification of key intermediates in the NAD⁺ salvage pathway. As expected, methylnicotinamide was significantly reduced in Derlin-2 KO cells (**Fig. 4C**). Additional intermediates, including NAD⁺ and nicotinamide mononucleotide, showed decreasing trends in both Derlin-2 and Derlin-3 KO cells, although they did not reach statistical significance. Altogether, loss of Derlin-2 is strongly associated with mitochondrial dysfunction (impaired oxidative respiration, attenuated mitochondrial Ca²⁺ efflux, and decrease in key metabolites in the NAD⁺ salvage pathway), whereas loss of Derlin-3 produces more modest effects, with a subset of mitochondrial defects overlapping those observed in Derlin-2 KO cells.

**Figure 4.**
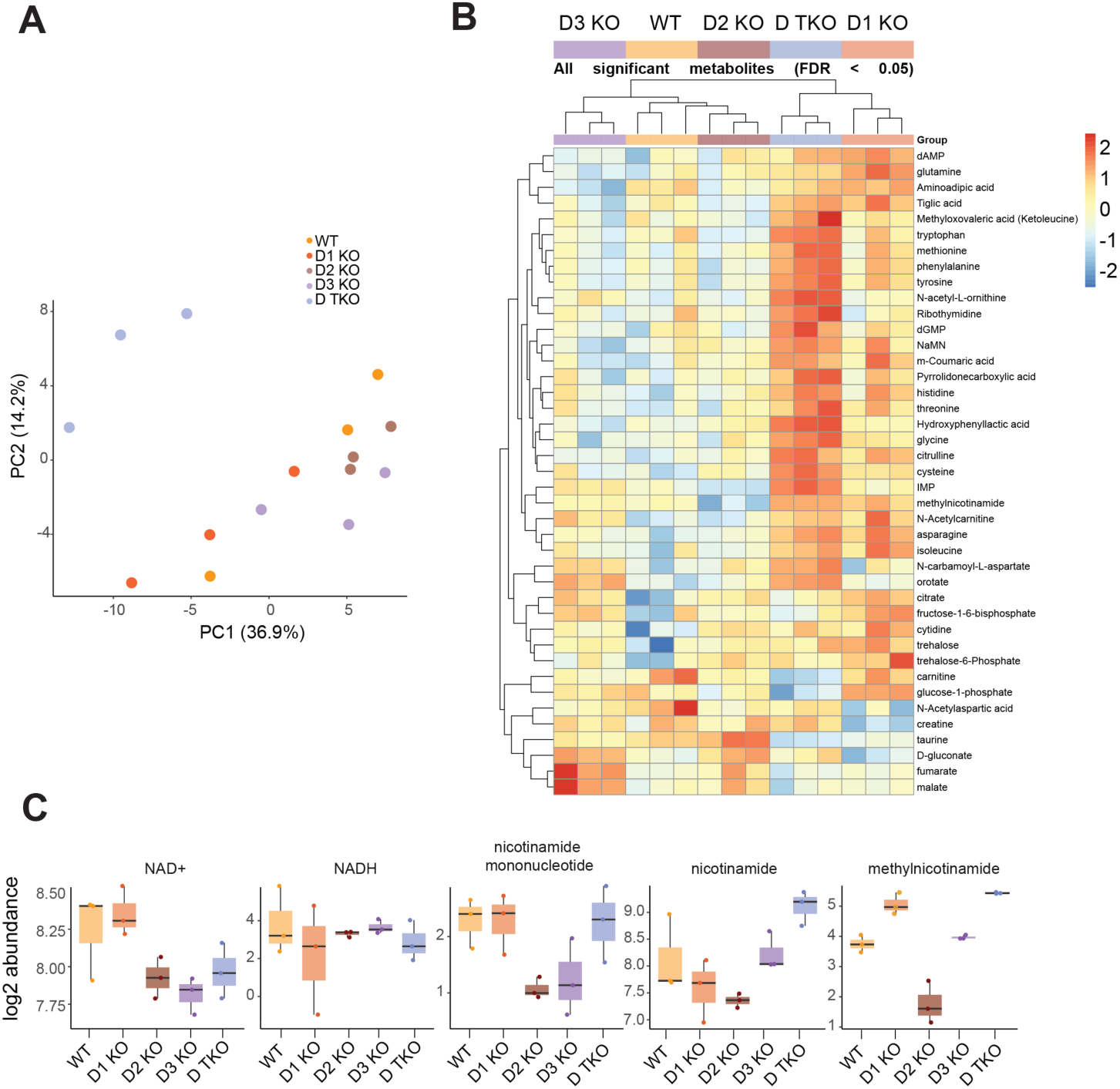
Loss of each Derlins lead to distinct metabolic changes. **(A)** Principal component analysis (PCA) was performed on log₂-transformed metabolite abundances from WT, Derlin-1 KO, Derlin-2 KO, Derlin-3 KO, and Derlin TKO HEK293 cells after removal of zero-variance metabolites. Data were centered and scaled, and samples are shown along the first two principal components, colored by their respective genotypes. **(B)** Heatmaps of significantly differentially abundant metabolites from WT, Derlin-1 KO, Derlin-2 KO, Derlin-3 KO, and Derlin TKO HEK293 cells were generated from log₂-transformed, row-scaled (Z-scored) data after exclusion of zero-variance metabolites. Samples were ordered by experimental group and hierarchically clustered using Euclidean distance and complete linkage. Heatmaps are shown for all groups and for individual contrasts, with metabolites annotated by direction of change relative to WT where indicated. **(C)** Box plots of log₂-transformed abundances of NAD⁺ salvage pathway intermediates across WT and Derlin knockout cell lines (Derlin-1 KO, Derlin-2 KO, Derlin-3 KO, and Derlin TKO). Boxes indicate the median and interquartile range, with individual points representing biological replicates.

### Proteomic profiling reveals distinct and overlapping consequences of Derlin loss

Because Derlins function in the ERAD pathway to selectively remove ER clients for proteasomal degradation, we hypothesized that loss of Derlin-2 and Derlin-3 may stabilize substrates that influence ER–mitochondria communication and mitochondrial function, prompting us to perform unbiased proteomic analyses. We performed tandem mass tag (TMT)–based quantitative proteomics in HEK293 WT, Derlin-1 KO, Derlin-2 KO, Derlin-3 KO, and Derlin TKO cells (**Fig. 5A**). Principal component analysis further demonstrated clear separation between WT and KO proteomes, with Derlin-2 KO clustering distantly from WT, Derlin-1 KO and Derlin TKO cells (**Fig. 5B**). Strikingly, Derlin-1 KO and TKO displayed the greatest overlap, sharing over 62 significantly altered proteins, while Derlin-2 and Derlin-3 KOs showed limited overlap with Derlin-1 KO (**Fig. 5C**). Volcano plots identified numerous significantly altered proteins relative to WT, with the largest changes observed in Derlin-1 KO and Derlin TKO cells (**Fig. 5D**). Gene ontology (GO) enrichment analysis revealed distinct signatures for each knockout (**Fig. 5E**). Derlin-1 KO cells were enriched for pathways linked to *organelle and cytoskeleton organization, actin filament processes, and cell junction assembly*. Derlin-2 KO cells displayed a narrower signature, with enrichment for *ER lumen, extracellular exosome, and lipase binding*, pointing to targeted disruption of ER/secretory functions. Derlin-3 KO cells showed enrichment for more regulatory categories, including *protein polymerization, microtubule nucleation, and cell growth regulation*, in line with its subtler cellular effects. Finally, Derlin TKO exhibited broad enrichment across *protein folding, ER stress responses, Golgi organization, trafficking, and metabolic processes*, reflecting the combined loss of all three homologs (**Fig. 5E**).

**Figure 5.**
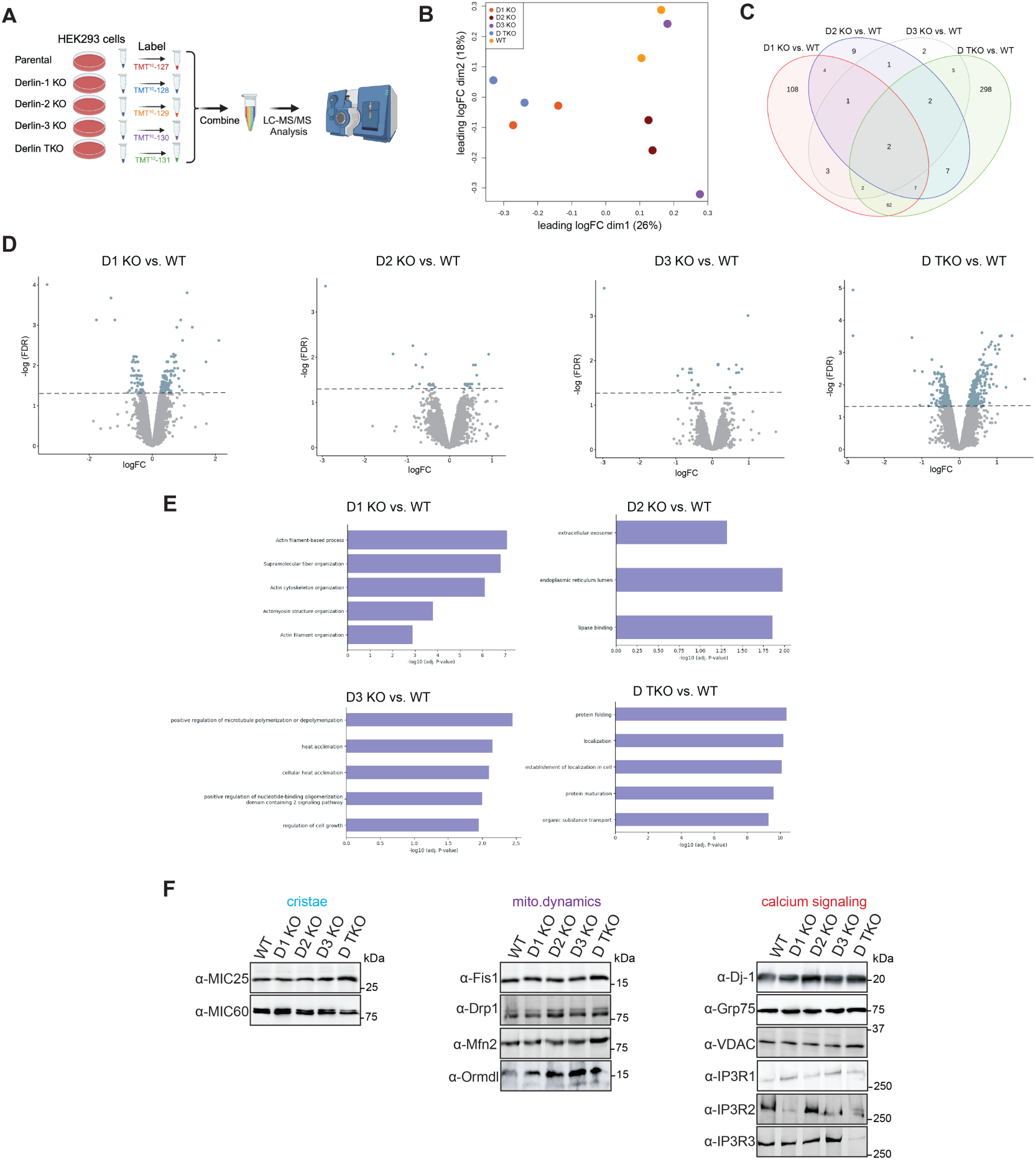
Proteomic analysis reveal ORMDLs are upregulated in Derlin-2 KO and Derlin-3 KO cells. **(A)** Experimental workflow of tandem mass tag (TMT) labeling coupled with LC–MS/MS for proteomic profiling of WT, Derlin-1 KO, Derlin-2 KO, Derlin-3 KO, and Derlin TKO HEK293 cell lines. **(B)** Multi-dimensional scaling (MDS) plot of the triplicate WT, Derlin-1 KO, Derlin-2 KO, Derlin-3 KO, and Derlin TKO samples. The x-axis shows the separation of the samples on the first dimension and accounts for 26% of the variability in the data, while the y-axis shows the second dimension and accounts for 28% of the variability. **(C)** Venn diagram of the significantly differentially abundant proteins in comparison with the WT samples with each Derlin KO cell line. **(D)** Volcano plots of the differentially abundant proteins in the comparison of each Derlin knockout with WT. For each plot, the x-axis represents the log fold change (logFC) with positive values increased in the knockout and negative values decreased in the knockout. The y-axis represents the negative log of the false discovery rate (-log (FDR)) with higher values corresponding to adjusted p-values closer to 0 (i.e. greater significance). Significantly differentially abundant proteins (adjusted p-value < 0.05) are represented by blue dots, while all other proteins are represented by gray dots. **(E)** Gene ontology (GO) enrichment analysis of differentially expressed proteins from (C), with representative terms shown according to adjusted significance values. **(F)** Validation of the steady-state levels of cristae, mitochondrial dynamics, and calcium signaling proteins by immunoblotting in WT, Derlin-1 KO, Derlin-2 KO, Derlin-3 KO, and Derlin TKO HEK293 cells. Data are representative of *n* = 3 independent experiments.

Given the pronounced effects of Derlin-2 and Derlin-3 loss on mitochondrial function, we hypothesized that these Derlins target substrates essential for mitochondrial function, particularly components of MERCs, where ER and mitochondrial proteins are in direct physical contact. Notably, proteomic analyses and immunoblotting revealed no significant changes in the abundance of core MERC components (DJ-1, GRP75, VDAC, IP3R1, mitochondrial dynamics regulators (FIS1, DRP1, MFN2), or even cristae-shaping factors (MIC25 and MIC60) (**Fig. 5F**). IP3R2 and IP3R3 protein levels were reduced in Derlin TKO cells compared to WT; however, RNA-seq analysis revealed that this decrease appears to be attributed to transcriptional downregulation rather than post-transcriptional regulation (**Fig. S4C,D**). Notably, ORMDL3 exhibited increased protein levels in Derlin-2 KO cells in comparison to WT (**Fig. 5D**). ORMDLs comprise three highly conserved mammalian homologs (ORMDL1–3) that negatively regulate sphingolipid biosynthesis by inhibiting the rate-limiting enzyme serine palmitoyltransferase (SPTLC)(Davis et al., 2018b). Consistent with this, we previously demonstrated that the yeast Derlin homolog Dfm1 regulates the abundance of the yeast ORMDL homolog Orm2 (Bhaduri, Aguayo, et al., 2022). Moreover, ORMDL proteins have been implicated in non-canonical roles in regulating mitochondrial dynamics (Sharma et al., 2024). Indeed, consistent with our proteomic findings, immunoblotting confirmed elevated steady-state levels of endogenous ORMDLs in both Derlin-2 and Derlin-3 KO cells (**Fig. 5F**).

### Derlin-2 and Derlin-3 targets ORMDL3 via ERAD pathway

Endogenous ORMDL antibodies do not distinguish amongst ORMDL paralogs due to their ∼80% sequence identity (Mahawar et al., n.d.). Accordingly, to determine which ORMDL paralog(s) are regulated by Derlin-2 and Derlin-3, we ectopically expressed ORMDL1-V5, ORMDL2-V5, or ORMDL3-V5 in WT, Derlin-1 KO, Derlin-2 KO, Derlin-3 KO, or Derlin TKO cells. Western blot analysis showed that all three ORMDL proteins were stabilized in Derlin-2 KO and Derlin-3 KO cells, whereas Derlin-1 KO had no effect (**Fig. 6A**). Among the paralogs, ORMDL3 exhibited the highest steady-state levels. To determine whether Derlin-2 and Derlin-3 are required for ORMDL degradation, we performed cycloheximide (CHX) chase assays. ORMDL3 underwent rapid turnover in WT, Derlin-2 KO, and Derlin-3 KO cells whereas its degradation was stabilized in Derlin-2 KO and Derlin-3 KO cells (**Fig. 6B**). To define the degradation pathway, WT, Derlin-2 KO, and Derlin-3 KO cells were transfected with ORMDL3-V5 and treated with inhibitors of the proteasome (MG132), lysosome (bafilomycin A1), or p97/VCP (CB5083) (**Fig. 6C**). As expected, ORMDL3 steady-state levels were elevated in Derlin-2 KO and Derlin-3 KO cells, consistent with impaired degradation. In agreement with previous reports that ORMDLs can undergo lysosomal-autophagic degradation (Wang et al., 2015), bafilomycin treatment increased ORMDL3 levels (**Fig. 6D**). Notably, in WT cells, inhibition of the proteasome or p97 markedly increased ORMDL3 steady-state levels (**Fig. 6D**), supporting a role for canonical ERAD in ORMDL3 turnover. In contrast, although ORMDL1 and ORMDL2 protein levels were elevated in Derlin-2 KO and Derlin-3 KO cells, they did not exhibit rapid degradation in WT cells that was comparable to ORMDL3 (**Fig. S5A**). Moreover, their steady-state levels were only modestly affected by p97 or bafilomycin treatment (**Fig. S5B,C**). ORMDL mRNA levels were unchanged in Derlin TKO cells, further supporting a post-transcriptional mechanism governing ORMDL abundance (**Fig. S4C,D**). Finally, we assessed whether Derlins influence ORMDL3 ubiquitination. WT, Derlin-2 KO, and Derlin-3 KO cells were co-transfected with ORMDL3-V5 and HA-ubiquitin. Immunoprecipitation revealed basal ubiquitination of ORMDL3 in WT cells, which was further increased in Derlin-2 and Derlin-3 KO cells (**Fig. 6E**), indicating that loss of Derlin-2 or Derlin-3 leads to accumulation of ubiquitinated ORMDL3, consistent with impaired ERAD. Finally, co-immunoprecipitation analyses demonstrated that both Derlin-2 and Derlin-3 associate with ORMDL3, further confirming that Derlin-2 and Derlin-3 target ORMDL3 for ERAD (**Fig. 6F**). Together, these data demonstrate that ORMDL3 has a faster turnover rate than ORMDL1 and ORMDL2 and is protein levels are regulated through both ERAD and lysosomal pathways.

**Figure 6.**
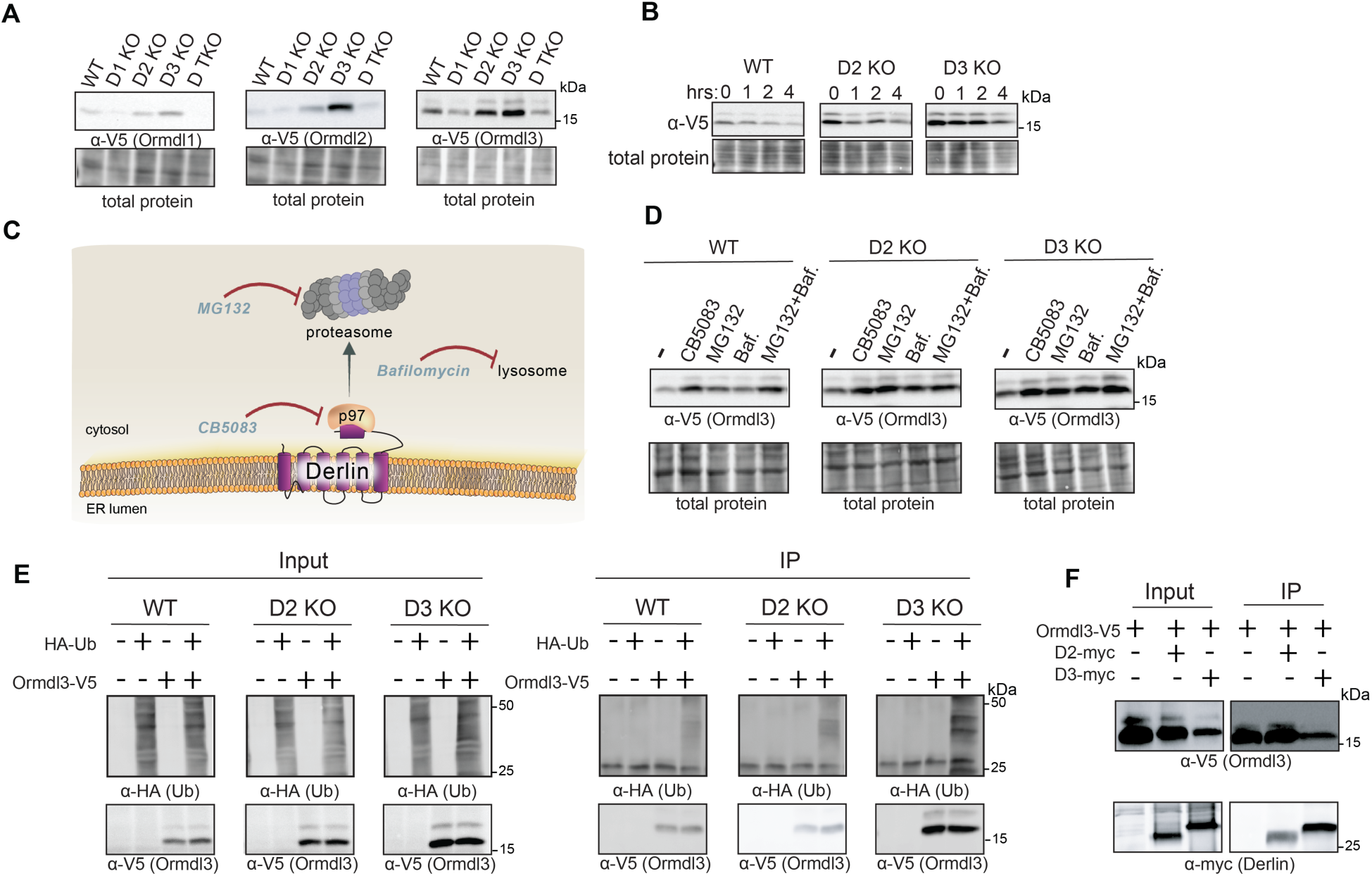
Derlin-2 and Derlin-3 regulate ORMDL ubiquitination and degradation. **(A)** WT, Derlin-1 KO, Derlin-2 KO, Derlin-3 KO, and Derlin TKO cells were transfected with ORMDL1-V5/His, ORMDL2-V5/His, or ORMDL3-V5/His for 24 h. Protein expression was analyzed by western blot. Data are representative of *n* = 3 independent experiments. **(B)** WT, Derlin-2 KO, and Derlin-3 KO HEK293 cells were transfected with ORMDL3-V5/His for 24 h, then treated with cycloheximide (10 µg/mL) for the indicated times up to 4 h. Protein stability was assessed by western blot. Data are representative of *n* = 3 independent experiments. **(C)** Schematic illustrating the autophagy and ERAD pathways, highlighting Derlins, p97, and the proteasome, and indicating the points of inhibition by CB5083 (p97), MG132 (proteasome), and bafilomycin A1 (lysosome). **(D)** WT, Derlin-2 KO, and Derlin-3 KO cells were transfected with ORMDL3-V5/His for 24 h, then treated for 6 h with vehicle, the p97/VCP inhibitor CB5083, the proteasome inhibitor MG132, or the lysosomal inhibitor bafilomycin. Protein levels were analyzed by western blot. Data are representative of *n* = 3 independent experiments. **(E)** HEK293 WT, Derlin-2 KO, and Derlin-3 KO cells were co-transfected with ORMDL3-V5/His and HA-ubiquitin for 24 h, followed by treatment with MG132 for 4 h. Lysates were immunoprecipitated with anti-V5 antibody, and blots were probed with anti-His, anti-HA, and anti-V5 antibodies. Data are representative of *n* = 3 independent experiments and shown as mean ± SEM. **(F)** HEK293 WT cells were transfected with ORMDL3-V5 and EV-Myc, Derlin-2-Myc, or Derlin-3-Myc followed by MG132 for 2 h, before pulling down on ORMDL3-V5 and analyzing the binding of Derlin-2-Myc and Derlin-3-Myc (3 biological replicates; *n* = 3).

### Derlin-dependent stabilization of ORMDL3 accumulates at MERCs

Amongst the ORMDL homologs, ORMDL3 has been identified as a genetic risk factor for childhood asthma, with single-nucleotide polymorphisms (SNPs) near the ORMDL3 locus associated with elevated expression levels (Acevedo et al., 2014). Beyond asthma, ORMDL3 variants have also been linked to other diseases, including inflammatory bowel disease, type 1 diabetes, atherosclerosis, multiple sclerosis, and certain cancers, highlighting its broader role in human health and disease(Ma et al., 2015; McGovern et al., 2010; Qiu et al., 2013; Zeng et al., 2025).

Recent studies in mice have begun to shed light on the pathological consequences of having elevated ORMDLs. For example, during inflammatory stimulation, ORMDL3 expression is upregulated, leading to its accumulation at MERCs. This delocalization has deleterious consequences, leading to impaired mitochondrial function and altered cellular homeostasis (Sharma et al., 2024). These findings suggest that not only the abundance but also the spatial distribution of ORMDLs is a key determinant of their pathogenic effects. Because we observed that all three ORMDL homologs, ORMDL1, ORMDL2, and ORMDL3, are stabilized in Derlin-2 and Derlin-3 KO cells, we hypothesized that elevated levels of these proteins could similarly promote their accumulation at MERCs. To test this, we performed biochemical fractionation (post-nuclear supernatant, microsomes, mitochondria, and MERCs fractions) followed by immunoblotting for all three ORMDLs, alongside markers of compartment identity: Sec61β for the ER/microsomes, TOMM20 for mitochondria, and SIGMA1R for MERCs.

In WT cells, all ORMDLs localized predominantly to the ER, consistent with prior reports. Strikingly, however, in Derlin-2 and Derlin-3 KO cells, only ORMDL3, but not ORMDL1 or ORMDL2, was selectively enriched at MERCs, despite all three homologs being stabilized (**Fig. 7A,B**). This selective delocalization of ORMDL3 points to a unique vulnerability of this paralog upon loss of Derlin function. To determine whether elevated ORMDL3 levels are sufficient to drive its accumulation at MERCs, we reintroduced Derlin-2 into the Derlin-2 KO background. Derlin-2 addback reduced overall ORMDL3 abundance and shifted its distribution back toward the bulk ER, markedly decreasing its enrichment at MERCs (**Fig. 7C,D).** To further determine whether Derlins limit ORMDL3 accumulation at ER–mitochondria contact sites, we performed proximity ligation assays (PLA) to quantify ORMDL3 enrichment at MERCs using antibodies against V5 for ORMDL3-V5 and the MERC marker VDAC1. ORMDL3–VDAC1 PLA signal was significantly increased in Derlin-2 KO cells relative to WT, and addback of Derlin-2 restored the PLA signal to near-WT levels (**Fig. 7E,F**). Altogether, we demonstrate that Derlins prevent the accumulation and delocalization of ORMDL3 to MERCs.

**Figure 7.**
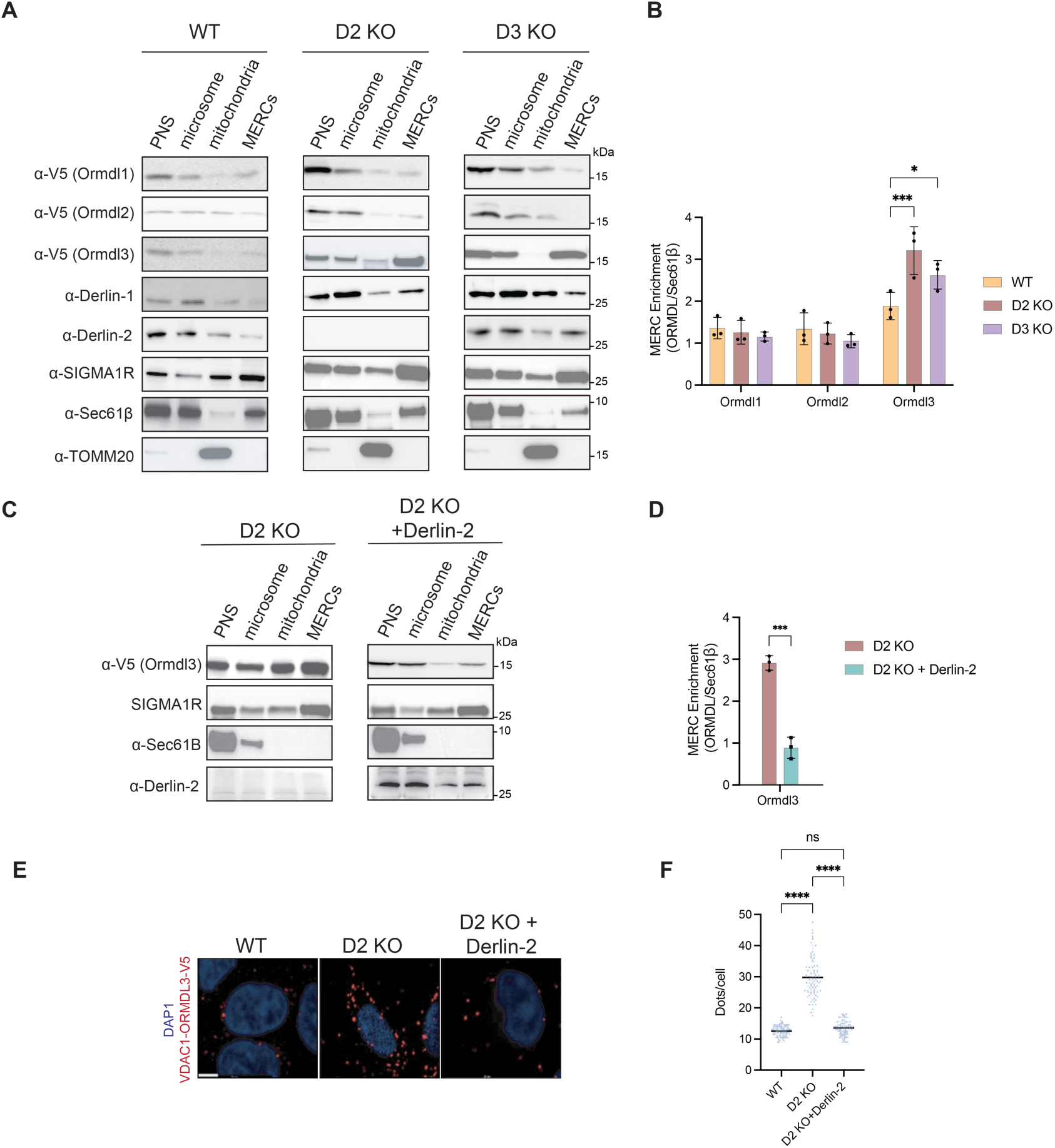
Derlin-2 and Derlin-3 limit ORMDL3 enrichment at mitochondria–ER contact sites. **(A)** Post-nuclear supernatant (PNS), mitochondria, endoplasmic reticulum (ER), and MERCs fractions were isolated from WT, Derlin-2 KO, and Derlin-3 KO cells expressing V5-tagged ORMDL1, ORMDL2, or ORMDL3. Immunoblot analysis was performed using anti-V5 antibodies to detect ORMDL proteins across subcellular fractions. Endogenous Derlin-1 and Derlin-2 were probed, and fraction purity and enrichment were validated using Sec61β as an ER marker, TOMM20 as a mitochondrial marker, and Sigma-1 receptor (Sigma1R) as a MERC marker. **(B)** MERC enrichment of V5-tagged ORMDL1, ORMDL2, and ORMDL3 was quantified from subcellular fractionation experiments in (A) ORMDL abundance in the MERC fraction was normalized to the ER marker Sec61B (ORMDL/Sec61) to account for ER content within contact sites. Data are shown as mean ± SEM from independent experiments, with individual data points representing biological replicates. Statistical significance was determined by one-way ANOVA with post-hoc multiple-comparison testing. *P < 0.05; ***P < 0.001.**(C-D)** MERC fractionation and quantification were performed as in (A) and (B), except that ORMDL distribution was analyzed in Derlin-2 KO cells with empty vector and Derlin-2 KO with ectopic expression of Derlin-2-Myc. **(E)** Images of *in situ* PLA (indicated in red) monitoring VDAC1-ORMDL3-V5 interaction in WT, Derlin-2 KO, and Derlin-2 KO with ectopic Derlin-2 cells. Scale bar, 10 uM. **(F)** Quantitative analysis of VDAC1-ORMDL3-V5 signals in WT, Derlin-2 KO, and Derlin-2 KO with ectopic Derlin-2 cells. Statistical significance was determined by one-way ANOVA followed by Tukey’s multiple-comparison test. *P < 0.05, **P < 0.01, ****P < 0.0001; ns, not significant.

### ORMDL3 Accumulation and Delocalization to MERCs Drives Mitochondrial Dysfunction

Given the pronounced mitochondrial defects observed in Derlin-2 KO cells, we focused all subsequent rescue experiments on this background. Collectively, Derlin-2 KO cells, and to a lesser extent Derlin-3 KO cells, exhibit overlapping mitochondrial phenotypes, including reduced MERC distance, mitochondrial fragmentation, diminished NAD⁺ salvage intermediates, impaired mitochondrial respiration, and attenuated Ca²⁺ efflux. We showed that ectopic expression of Derlin-2 in Derlin-2 KO cells not only reduced ORMDL3 steady-state levels but also promoted relocalization of ORMDL3 from MERCs back to the ER (**Fig. 7C–F**). We therefore hypothesized that restoring ORMDL3 localization in the ER would rescue mitochondrial function. Consistent with this, ectopic expression of Derlin-2 in Derlin-2 KO cells significantly increased basal, maximal, and spare respiratory capacity to near WT levels (**Fig. 8A,B**). If elevated ORMDL3 levels drive its aberrant enrichment at MERCs, then reducing endogenous ORMDL3 in Derlin-2 KO cells should similarly restore mitochondrial respiration. To test this, we generated siRNAs targeting each ORMDL paralog to selectively reduce their expression (**Fig. 8C; Fig. S6A,B**) and assessed the resulting effects on mitochondrial function. Remarkably, knockdown of ORMDL3, but not ORMDL1 or ORMDL2, in Derlin-2 KO cells restored mitochondrial respiration to near WT levels (**Fig. 8D; Fig. S6C,D**). Notably, individual knockdown of ORMDL1, ORMDL2, or ORMDL3 in WT cells did not alter mitochondrial respiration (**Fig. 8D; Fig. S6C,D**), underscoring a specific requirement for Derlin-2–dependent regulation of ORMDL3 stability in maintaining mitochondrial function. Moreover, selective knockdown of ORMDL3 in Derlin-2 KO cells rescued the rate of mitochondrial Ca²⁺ efflux, further indicating that ORMDL3 accumulation is a primary driver of the mitochondrial defects observed in Derlin-2 KO cells (**Fig. 8E**).

**Figure 8.**
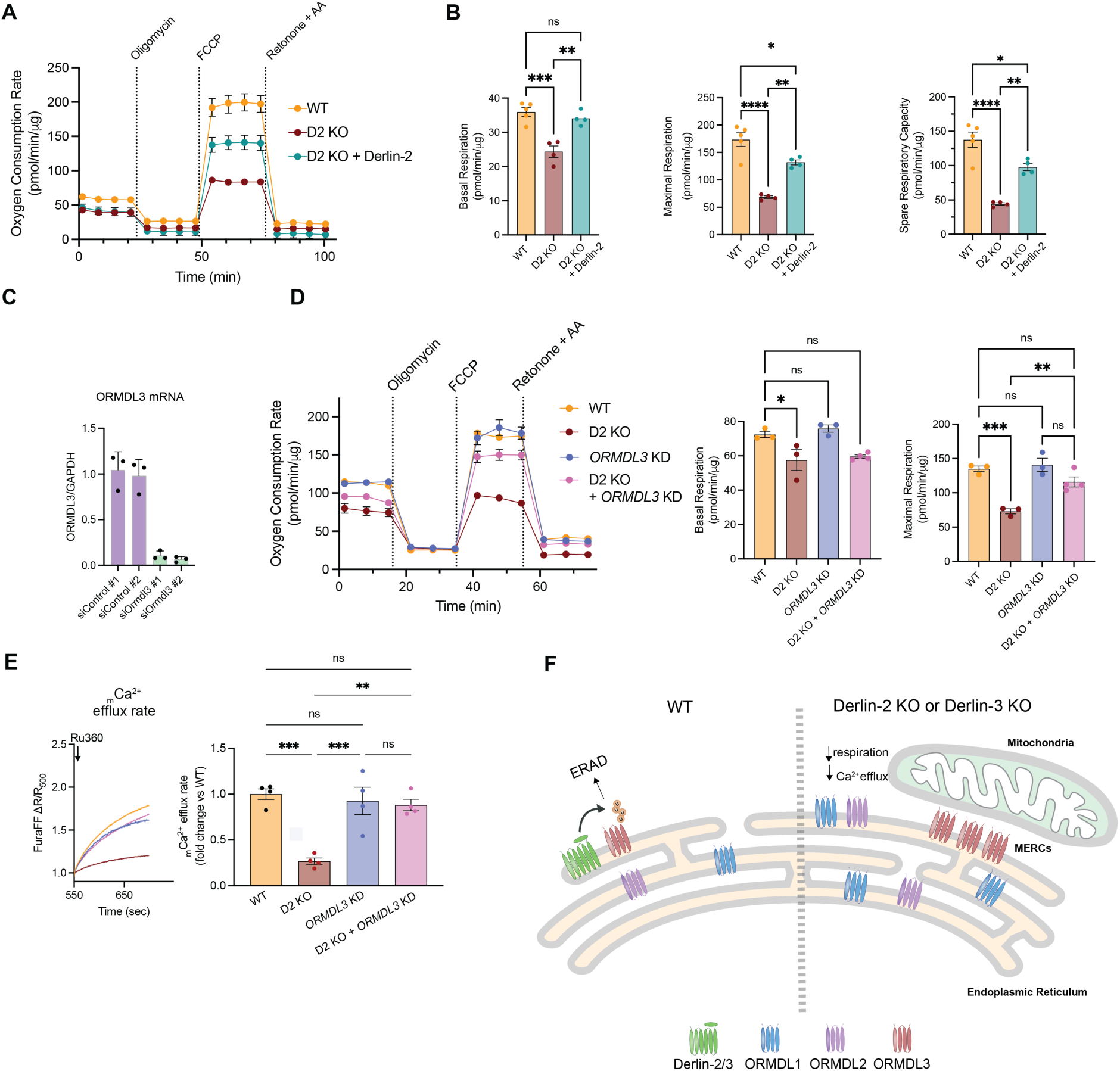
Derlin-2 regulates mitochondrial respiration and Ca²⁺ homeostasis through ORMDL3. **(A)** Oxygen consumption rate (OCR) was measured in HEK293 WT, Derlin-2 KO, and Derlin-2 KO with ectopic expression of Derlin-2 using a Seahorse Analyzer. **(B)** Quantification of basal OCR, maximal respiration, and spare respiratory capacity were quantified from (A). Data represent *n* = 4 independent experiments (mean ± SEM). Statistical significance was determined by one-way ANOVA followed by Tukey’s multiple-comparison test. *P < 0.05, **P < 0.01, ***P < 0.001, ****P < 0.0001; ns, not significant. **(C)** ORMDL3 mRNA levels were measured by quantitative RT–PCR in cells transfected with two independent control siRNAs (siControl #1 and #2) or two independent ORMDL3-targeting siRNAs (siORMDL3 #1 and #2). ORMDL3 expression was normalized to GAPDH and is shown relative to control conditions. Data are presented as mean ± SEM from independent experiments, with individual data points representing biological replicates. **(D)** Oxygen consumption rate was measured by Seahorse extracellular flux analysis in WT, Derlin-2 KO, ORMDL3 KD, and Derlin-2 KO with ORMDL3 KD HEK293 cells. Basal and maximal respiration were quantified following sequential injection of oligomycin, FCCP, and rotenone/antimycin A. Data are shown as mean ± SEM from independent experiments, with individual data points representing biological replicates. Statistical significance was determined by one-way ANOVA followed by Tukey’s multiple-comparison test. *P < 0.05, **P < 0.01, ***P < 0.001; ns, not significant. **(E)** Ratiometric monitoring of mitochondrial Ca²⁺ dynamics in WT, Derlin-2 KO, ORMDL3 KD, and Derlin-2 KO with ORMDL3 KD HEK293 cells using FuraFF. Mitochondrial Ca²⁺ (mCa²⁺) efflux rate was normalized at 550 sec. Fold change in mCa²⁺ efflux of Derlin KO cells was compared to WT. Data represent *n* = 4 independent experiments (mean ± SEM). Statistical significance was determined by one-way ANOVA followed by Tukey’s multiple-comparison test. *P < 0.05, **P < 0.01; ns, not significant. **(F)** Schematic illustrating how Derlin-2 and Derlin-3 controls ORMDL3 distribution between the ER and MERCs. In WT cells, Derlin-2 and Derlin-3 promotes ERAD of all ORMDLs, particularly restricting ORMDl3 accumulation at MERCs and thereby maintaining proper ER–mitochondria communication, calcium transfer, and mitochondrial respiratory function. In the absence of Derlin-2 or Derlin-3, ORMDL3 aberrantly accumulates at MERCs, leading to disrupted ER–mitochondria signaling and mitochondrial dysfunction. This model integrates the observed changes in ORMDL3 localization, MERC architecture, and mitochondrial physiology in Derlin-2–deficient and to some extent Derlin-3-deficient cells.

## Limitations of study

Localization conclusions depend on overexpressed, epitope-tagged ORMDL paralogs rather than endogenous proteins. The field lacks commercial antibodies for endogenous ORMDL detection, forcing reliance on tagged overexpression from 2007 to present. To get around this, we combined orthogonal approaches such as MERC fractionation, PLA, Derlin-2 rescue, and knockdown of endogenous ORMDL3 to support ORMDL3 MERC enrichment studies.

## Discussion

We identify previously unrecognized roles for Derlins in safeguarding mitochondrial homeostasis through selective ERAD of ORMDL proteins. Our data indicate that Derlin-2 and Derlin-3 mainly target ORMDL3 for ERAD. Although loss of Derlin-2 or Derlin-3 leads to stabilization of all three ORMDL proteins at the steady-state level, only increased levels of ORMDL3 results in aberrant delocalization to MERCs, leading to impaired mitochondrial respiration and defective mitochondrial Ca^2+^ handling (**Fig. 8F**). Reducing ORMDL3 levels in a Derlin-2 KO and *not* WT background restored mitochondrial function in comparison to Derlin-2 KOs, whereas depletion of ORMDL1 or ORMDL2 had no effect, establishing ORMDL3 as the pathogenic isoform in this context. These findings uncover Derlin-dependent ERAD limits ORMDL3 increased proteins levels and mis-localization to MERCs in order to safeguard mitochondria mitochondria. Importantly, this work provides the first evidence that Derlin-dependent ERAD functions not only in the clearance of misfolded or orphan proteins, but also in regulating protein abundance to ensure proper localization of client proteins such as ORMDL3, an evolutionarily conserved ER membrane protein with broad implications in human disease, including inflammation, cancer, asthma, inflammatory bowel disease, type 1 and type 2 diabetes, multiple sclerosis, obesity, and nonalcoholic fatty liver disease(Brown & Spiegel, 2023; Davis et al., 2018c, 2018d; H. Wang et al., 2025).

Herein, we also present the first systematic characterization of all three Derlin paralogs and delineate several foundational differences among them. Notably, loss of Derlin-2 uniquely activates UPR, whereas Derlin-1 and Derlin-3 KO cells do not, indicating that Derlin-2 serves as the principal paralog responsible for alleviating ER stress. Metabolomic and proteomic profiling further reinforced the differences amongst Derlin paralogs. The metabolomic landscape differed markedly across knockout lines: Derlin-1 KO cells exhibited the most extensive metabolic remodeling, with prominent enrichment in primary bile acid biosynthesis, amino acid metabolism, taurine and hypotaurine metabolism, thiamine metabolism, and porphyrin metabolism. In contrast, Derlin-2 and Derlin-3 KOs showed fewer global metabolic changes but shared a selective disruption of the NAD⁺ salvage pathway, consistent with their effects on mitochondrial respiration and bioenergetics. Proteomic analyses mirrored these trends. Derlin-1 KO cells displayed the broadest proteomic alterations, with differential expression of proteins involved in organelle organization, cytoskeletal architecture, actin filament dynamics, and cell–cell junction assembly. By contrast, Derlin-2 KO cells exhibited a more focused proteomic signature enriched for ER lumen, extracellular exosome, and lipase-binding categories, suggesting targeted impairment of ER and secretory pathway functions. Derlin-3 KO cells showed the fewest proteomic changes. Notably, combined loss of all three Derlins resulted in upregulation of canonical UPR components, underscoring partial functional redundancy among Derlins in maintaining ER proteostasis.

Although ORMDL1–3 share near-identical transmembrane topology and ER localization, our findings reveal isoform-specific functional divergence. Specifically, ORMDL3, but not ORMDL1 or ORMDL2, redistributes to MERCs upon loss of Derlin-2 or Derlin-3. This spatial delocalization appears to be a critical determinant of mitochondrial dysfunction. Whereas accumulated ORMDL1 and ORMDL2 remain confined to the bulk ER, ORMDL3 preferentially engages the MERC interface, coinciding with downstream mitochondrial defects, including impaired mitochondrial respiration and defective mitochondrial Ca²⁺ efflux. Notably, a recent study reported that under inflammatory conditions, elevated ER-resident ORMDL3 similarly delocalizes to MERCs (Sharma et al., 2024). Together, these observations suggest that increased ORMDL3 levels promote its recruitment and selective sequestration of other factors at MERCs. This raises an intriguing question: despite high sequence similarity among ORMDL paralogs, what molecular features uniquely enable ORMDL3 to engage MERCs and recruit additional components? Addressing this question represents an important avenue for future investigation. Moreover, to date, no studies have delineated pathological differences among the individual ORMDL homologs. Our findings provide mechanistic insight into why ORMDL3 emerges as the pathogenic isoform, whose elevated levels contribute to a broad spectrum of diseases.

Emerging evidence places sphingolipids at the intersection of ER proteostasis, mitochondrial function, and inflammation. Multiple studies indicate that localized sphingolipid dysregulation at MERCs can destabilize mitochondrial structure and signaling (Fugio et al., 2020). Reduced sphingolipid synthesis has been shown to alter membrane fluidity and curvature, disrupt Ca²⁺ channel organization at MERCs, and impair mitochondrial respiration(Planas-Serra et al., 2023). Indeed, when we performed bulk lipidomic analyses, Derlin-2 KO cells exhibited selective reductions in specific sphingolipid species (**Fig. S7**). If reduced sphingolipid biosynthesis at MERCs were the primary driver of mitochondrial dysfunction, pharmacologic suppression of serine palmitoyltransferase using myriocin would be expected to phenocopy the defects observed in Derlin-2 KO cells. In contrast, several studies have shown that global inhibition of sphingolipid synthesis with myriocin induces elongated, hyperfused mitochondria without compromising mitochondrial respiration or Ca²⁺ signaling, effects opposite to the mitochondrial defects we observed in Derlin-2 KO cells(Ebert et al., 2025; Smith et al., 2013). These findings suggest that mitochondrial dysfunction is unlikely to arise solely from ORMDL3’s canonical role in suppressing sphingolipid biosynthesis. Consistent with this interpretation, prior studies have shown that aberrant accumulation of ORMDL3 at MERCs, both *in vitro* and in inflammatory disease models in mice, promotes the recruitment of inflammatory signaling factors to these interfaces (Sharma et al., 2024). Moreover, our work and that of others demonstrates that ORMDL3 mislocalization enhances ER–mitochondria tethering and drives mitochondrial fragmentation, supporting a model in which aberrant ORMDL3 localization, and potential sequestration of associated factors, disrupts mitochondrial function (Sharma et al., 2024). Notably, the earlier investigation focused primarily on ORMDL3 and did not systematically evaluate the functional contributions of the closely related paralogs ORMDL1 and ORMDL2. In contrast, our analysis directly compares all three ORMDL family members, revealing paralog-specific differences in localization and regulation by Derlins. Defining how selective ORMDL enrichment at MERCs perturbs mitochondrial function will therefore be essential for understanding its contribution to disease pathogenesis, while also establishing a broader framework in which Derlins differentially regulate ORMDL paralogs to preserve organelle homeostasis.

Although both Derlin-2 and Derlin-3 KO cells exhibit pronounced mitochondrial dysfunction, the physiological consequences are substantially more severe in Derlin-2 KO cells. Loss of Derlin-2 results in markedly impaired mitochondrial respiration, defective Ca²⁺ efflux, and heightened ER stress, establishing Derlin-2 as the primary driver of the mitochondrial phenotypes, with partial functional redundancy provided by Derlin-3. Consistent with this model, Derlin-2 KO cells exhibit significantly greater ORMDL3 enrichment at MERCs compared with Derlin-3 KO cells, further establishing Derlin-2 as the dominant paralog regulating ORMDL3 localization. Perhaps most intriguingly, Derlin-1 KO cells also exhibit mitochondrial fragmentation but do not display measurable defects in mitochondrial function. In contrast to Derlin-2 and Derlin-3 knockout cells, Derlin-1 knockout cells exhibited a significant increase in ER area accompanied by an increased MERC distance. Notably, gene ontology analysis of the Derlin-1 knockout proteome revealed enrichment for pathways involved in organelle and cytoskeletal organization, actin filament dynamics, and cell junction assembly. These alterations may underlie the proliferative defects observed in Derlin-1 knockout cells and contribute to the broader disorganization of both mitochondrial and ER architecture. Given the extensive metabolomic, proteomic, and lipidomic changes associated with Derlin-1 loss, an important avenue for future investigation will be to define the mechanistic basis by which Derlin-1 regulates cellular homeostasis and structural organization.

In summary, this work establishes Derlin-dependent ERAD as a critical regulator of mitochondrial homeostasis by governing the abundance and spatial distribution of ORMDL3 at ER–mitochondria contact sites. Our findings reveal that Derlin-2, with partial redundancy from Derlin-3, selectively restrains ORMDL3 accumulation and prevents its aberrant enrichment at MERCs, thereby preserving mitochondrial respiration, Ca²⁺ flux, and bioenergetic integrity. Importantly, these data redefine ERAD not simply as a quality-control pathway for misfolded proteins, but as a mechanism for fine-tuning protein levels of ER clients whose mislocalization can profoundly disrupt organelle communication. By uncovering ORMDL3 as a key effector linking ER proteostasis to mitochondrial dysfunction, this study provides a conceptual framework for understanding how dysregulated ERAD contributes to metabolic and inflammatory disease states and highlights Derlins as potential therapeutic targets for disorders driven by ORMDL3 dysregulation.

## Acknowledgements

We thank Hideki Nishitoh (University of Miyazaki) for providing the Derlin KO cell lines and Dr. Majid Ghassemian of the Biomolecular and Proteomics Mass Spectrometry Facility for processing our samples for mass spectrometry. We would also like to acknowledge the Huck Institutes’ Metabolomics Core Facility (RRID:SCR_023864) for use of the OE 240 LCMS and Sergei Koshkin for helpful discussions on sample preparation and analysis. Finally, we thank the Neal lab members for in-depth discussions and technical assistance. These studies were supported by Howard Hughes Medical Institute Freeman Hrabowski Program, NIH grant 1R35GM133565-01, NSF CAREER grant 2047391, and CZI Science Diversity Leadership Award (to S.E.N.); CZI Science Diversity Leadership Award (to A.H.J.); Harold S. Geneen Charitable Trust Awards Program (to D.T.); Howard Hughes Medical Institute Hanna H. Gray Fellows Program Faculty Phase grant GT1565 and the Burroughs Welcome Fund PDEP Transition to Faculty grant 1022604 (to M.R.M); the NIH Grant T32GM154124 (to V.R.B); and American Australian Association (to N.A.S.).

## Author contributions

Designed research, N.A.S., J.M.A., A.G.M., J.C.S., A.A.K., C.S.C., V.R.B., P.P., D.T., M.R.M., A.H., and S.E.N.; performed research, N.A.S., J.M.A., A.G.M., J.C.S., A.A.K., C.S.C., V.R.B., D.T., and A.H.; analyzed data, N.A.S., J.M.A., A.G.M., J.C.S., A.A.K., C.S.C., V.R.B., P.P., M.A.P., D.T., M.R.M., A.H., and S.E.N.; wrote the paper, S.E.N. and N.A.S. All authors reviewed the results and approved the final version of the manuscript.

## Data availability

The metabolomics, lipidomics, proteomics, and RNA-Seq datasets generated during this study are available in the Mendeley Data repository at DOI: 10.17632/x7cdvppxj5.

## Materials & Methods

### Plasmids

The protein-coding sequences of ORMDL1, ORMDL2, ORMDL3, were ordered as gBlocks and inserted into the pcDNA3.1/V5-His vector (ThermoFisher) using NEBuilder HiFi DNA Assembly according to the manufacturer’s protocol to generate pcDNA3.1 ORMDL1–V5/His, pcDNA3.1 ORMDL2–V5/His, and pcDNA3.1 ORMDL3–V5/His. The protein-coding sequence of Derlin-1, Derlin-2, and Derlin-3 were ordered as gBlocks and inserted into the pcDNA3.1/Myc-His vector (ThermoFisher) using NEBuilder HiFi DNA Assembly according to the manufacturer’s protocol to generate pcDNA3.1 Derlin-1–Myc/His, pcDNA3.1 Derlin-2–Myc/His, and pcDNA3.1 Derlin-3–Myc/His. The pRK5 HA-Ubiquitin plasmid was acquired from Addgene. Plasmid sequences were verified with Sanger (Eton Bioscience) and/or whole plasmid (Plasmidsaurus) sequencing.

### TOPO cloning and sequencing

Genomic DNA from HEK293 WT, Derlin-3 KO, and Derlin Triple KO cells was extracted using Monarch Genomic DNA Purification Kit (NEB Labs) according to manufacturer’s instructions. The area over Exon 1 of Derlin-3, where CRISPR genetic editing occurred, was amplified using GoTaq Green Master Mix (Promega) with primers forward: GGCTGGGAGGCGGGTTAAAG and reverse: CAGTTTCCCCGTCCCGGC. Amplicons were verified on an agarose gel, then incorporated using the TOPO TA Cloning Kit for Sequencing (Invitrogen) according to manufacturer’s instructions. Plasmids were then transformed into Mix & Go Competent cells according to manufacturer’s instructions, grown overnight at 37°C on LB + Ampicillin plates. Colonies were picked and sent off for sequencing at Plasmidsaurus.

### Cell Culture

HEK293 cell wild type, HEK293 Derlin-1 KO, HEK293 Derlin-2 KO, HEK293 Derlin-3 KO, and HEK293 Derlin Triple KO were a generous gift from Professor Hideki Nishitoh, University of Miyazaki. Cells were grown in 10% (v/v) fetal bovine serum (FBS)/DMEM high glucose at 37°C 5% CO_2_. Cells were seeded, then transfected the following day if required, and/or treated as indicated in the Figure legends.

### Plasmid Transfections

For plasmid transfections, 300,000-600,000 cells were seeded into a 6-well plate, then transfected the following day with a total of 1 μg plasmid using 4 μL Lipofectamine 3000 reagent and 2 μL P3000 reagent (Invitrogen) for 24 h (unless indicated otherwise in the Figure Legends).

### Growth Curves

HEK293 WT and Derlin KO cells were seeded at 300 cells per well in a 96-well plate and allowed to adhere before imaging of cell confluence with the Incucyte SX5 every hour for 6 days. For treated cells, 200 ng/mL Tunicamycin or vehicle was added at the 0 h timepoint. Confluence was normalised to the 0 h time point for each cell line. To quantify growth over time, the area under each growth curve was measured and normalised to the WT, which was set to 1.

### RT-qPCR

Using TRIzol reagent (Invitrogen), total RNA was isolated from tissues and further purified with the rNeasy kit (Qiagen Inc). RNA concentration was determined by measuring absorbance at 260 nm and 280 nm using a NanoDrop 1000 spectrophotometer (NanoDrop products, Wilmington, DE, USA). Approximately 1 μg of RNA was reverse-transcribed using a High-Capacity cDNA Reverse Transcription Kit (Applied Biosciences, Carlsbad, CA). Quantitative PCR (qPCR) was then performed using SYBR Green (Life Technologies, Carlsbad, CA). For qPCR, 50 ng of cDNA was loaded into each well of a 384-well plate, with the reaction carried out on an ABI Prism 7900HT system (Applied Biosystems) with the following cycle: 1 cycle at 95°C for 10 min; 40 cycles of 95°C for 15 s; 59°C for 15 s, 72°C for 30 s, 800 and 78°C for 10 s; 1 cycle of 95°C for 15 s; 1 cycle of 60°C for 15 s; and one cycle of 95°C for 15 s. GAPDH normalization was used to present the data as fold changes.

### RNA-Seq

Samples for RNA-Seq were prepared according to Plasmidsaurus processing and analysis instructions. 200,000 HEK WT and Derlin Triple KO cells were isolated and washed 2 times with PBS before resuspending in DNA/RNA Shield Stabilization Reagent (Zymo Research) before being sent to Plasmidsaurus. Quality of the fastq files was assessed using FastQC v0.12.1. Reads were then quality filtered using fastp v0.24.0 with poly-X tail trimming, 3’ quality-based tail trimming, a minimum Phred quality score of 15, and a minimum length requirement of 50 bp. Quality-filtered reads were aligned to the reference genome using STAR aligner v2.7.11 with non-canonical splice junction removal and output of unmapped reads, followed by coordinate sorting using samtools v1.22.1. PCR and optical duplicates were removed using UMI-based deduplication with UMIcollapse v1.1.0. Alignment quality metrics, strand specificity, and read distribution across genomic features were assessed using RSeQC v5.0.4 and Qualimap v2.3, with results aggregated into a comprehensive quality control report using MultiQC v1.32. Gene-level expression quantification was performed using featureCounts (subread package v2.1.1) with strand-specific counting, multi-mapping read fractional assignment, exons and three prime UTR as the feature identifiers, and grouped by gene_id. Final gene counts were annotated with gene biotype and other metadata extracted from the reference GTF file. Sample-sample correlations for sample-sample heatmap and PCA were calculated on normalized counts (CPM) using Pearson correlation on ClustVis. Differential expression was done with edgeR v4.0.16 using standard practice including filtering for low-expressed genes with edgeR::filterByExprwith default values.

### Quantification of TEM Micrographs and Parameters Using ImageJ

Samples were fixed in a manner to avoid any bias, per established protocols(Ainbinder et al., 2023; Lam et al., 2021; Neikirk et al., 2023). Following preparation, cells were embedded in 100% Embed 812/Araldite resin with polymerization at 60 °C overnight. After ultrathin sections (90–100 nm) were collected, they were post-stained with lead citrate and imaged (JEOL 1400+ at 80 kV, equipped with a GatanOrius 832 camera). The National Institutes of Health (NIH) *ImageJ* software was used to quantify TEM images, as described previously

### Live-Cell Imaging and Analysis

Live-cell mitochondrial dynamics were visualized using a Nikon Eclipse Ti2 inverted fluorescence microscope equipped with a Yokogawa CSU-W1 spinning disk confocal scanner, Hamamatsu Fusion BT camera, SoRa super-resolution module, environmental chamber, piezo stage controller, and solid-state lasers (405, 488, 561, and 640 nm), all controlled via NIS-Elements AR software (version 5.42). Cells cultured in 35-mm MatTek glass-bottom dishes were imaged with a 100× Plan Apo Lambda D oil immersion objective (NA 1.45), capturing MitoTracker Orange-labeled mitochondria through the 561 nm channel at 2.5-second intervals over 5 minutes using either standard W1 mode (xy pixel size: 65 nm) or SoRa mode (xy pixel size: 23 nm), with laser power maintained at 5% and 100 ms exposure time per frame to balance phototoxicity concerns with signal quality (SNR ∼1.5-2.5). The Perfect Focus System maintained z-stability throughout all acquisitions. At the same time, high-resolution z-stacks (10-20 µm thick) were acquired in SoRa mode with 100 nm step sizes and subsequently enhanced using the Nikon Batch Deconvolution module (version 6.10.02), implementing Blind and Richardson-Lucy algorithms with 20 iterations and automatic noise estimation. SORA imaging and image processing were performed, in part, through the Vanderbilt Cell Imaging Shared Resource and Nikon Center of Excellence (supported by NIH grants CA68485, DK20593, DK58404, DK59637, and EY08126).

### Pharmacologic Inhibitor Treatments for Confocal Microscopy

For confocal imaging experiments, cells were treated with selective small-molecule inhibitors targeting key mitochondrial and ER-associated pathways. All inhibitor treatments were performed under identical culture conditions, with concentrations and incubation times optimized to induce acute pathway perturbation while minimizing secondary cytotoxicity. Cells were treated with an OPA1 inhibitor at 50 µM for 4–6 hours to disrupt inner mitochondrial membrane fusion and cristae maintenance. MFN2 activity was inhibited with a compound at 20 µM for 4–6 hours, targeting outer mitochondrial membrane fusion and mitochondria–ER tethering. To acutely perturb cristae organization, cells were treated with a MICOS complex inhibitor at 30 µM for 20 minutes, rapidly disrupting cristae junction stability. ER-associated degradation pathways were inhibited with Derlin inhibitors targeting Derlin-1, -2, and -3, as well as a pan-Derlin triple-target strategy, applied at 20 µM for 4–6 hours. Following treatment, cells were immediately processed for live or fixed confocal imaging as indicated, using identical acquisition settings across all experimental conditions.

### siRNA and cell KOs for TEM, Oxygen Consumption, and Ca+2 Flux Assays

siRNA reagents targeting mitochondrial dynamics, cristae organizing, and ORMDL genes were obtained from Thermo Fisher Scientific and used for knockdown experiments in HEK293 cells. Exon specific siRNAs were directed against the following human gene targets: MFN2 mitofusin 2 Entrez Gene ID 9927 located on chromosome 1 positions 11980181 to 12013515 on build GRCh38 also known as CMT2A CMT2A2 CPRP1 HMSN6A HSG and MARF using siRNA ID 138495; OPA1 mitochondrial dynamin like GTPase Entrez Gene ID 4976 located on chromosome 3 positions 193593144 to 193697811 on build GRCh38 also known as BERHS MGM1 MTDPS14 NPG NTG and largeG using siRNA ID 144409; IMMT inner membrane mitochondrial protein also referred to as MIC60 Entrez Gene ID 10989 located on chromosome 2 positions 86143932 to 86195770 on build GRCh38 also known as HMP MINOS2 Mic60 P87 P88 89 P89 PIG4 and PIG52 using siRNA ID 136127; CHCHD3 coiled coil helix coiled coil helix domain containing 3 Entrez Gene ID 54927 located on chromosome 7 positions 132784862 to 133082158 on build GRCh38 also known as MINOS3 Mic19 and PPP1R22 using siRNA ID 147671; CHCHD6 coiled coil helix coiled coil helix domain containing 6 Entrez Gene ID 84303 located on chromosome 3 positions 126704220 to 126960420 on build GRCh38 also known as CHCM1 Mic25 and PPP1R23 using siRNA ID 182681; ORMDL3 using siRNA ID 228503; ORMDL1 using siRNA ID 41257; and ORMDL2 using siRNA ID 26474. All siRNAs were designed against Homo sapiens transcripts and were transfected at a final concentration of 10 nM using Lipofectamine RNAiMAX Thermo Fisher Scientific according to the manufacturer’s instructions.

### Mitochondrial Volume, Surface Area, and Morphology

Mitochondrial networks were quantified by measuring network volume in Imaris (Bitplane, Concord, MA). W1 and SORA images were acquired as a z-stack and reconstructed into a 3D volume in Imaris. Voxel dimensions and z-step size were preserved during image import to ensure accurate spatial scaling across all samples. The resulting images were converted to surface renderings. Imaris identifies individual mitochondria and interconnected networks as surfaces.

To automate image analysis, parameters for surface rendering, including intensity thresholding, background subtraction, and minimum object size, were set at the start of the study for each dataset and applied identically across all experimental conditions. The remaining samples were rendered in batches to ensure consistency and minimize user bias.

Once the surfaces were rendered, mitochondrial volume, surface area, and sphericity were quantified for each surface. All analyses were performed using uniform segmentation and rendering parameters across conditions, and quantification was conducted in a blinded manner when applicable to reduce bias further.

### Data Analysis

GraphPad Prism (La Jolla, CA, USA) was used for all statistical analyses. All experiments involving confocal and TEM data included at least three independent replicates. The experimenters did not handle statistics. The black bars represent the standard error of the mean. For all analyses, one-way ANOVA was performed with tests against each independent group, and significance was assessed using Fisher’s protected least significant difference (LSD) test. *, **, ***, **** were set to show significant difference, denoting *p* < 0.05, *p* < 0.01, *p* < 0.001, and *p* < 0.0001, respectively.

### *In vivo* ubiquitination pulldown

3.5×10^6^ HEK293, HEK293 Derlin-2 KO, and HEK293 Derlin-3 KO cells were seeded in a T-75 flask and transfected the following day with a total of 6 μg plasmid (at a ratio of 2.5 μg pcDNA3.1 ORMDL3-V5-His : 1 μg pRK5 HA-Ubiquitin, and pcDNA3.1 EV-V5 as needed) using 24 μL Lipofectamine 3000 reagent and 12 μL P3000 supplemental reagent (Invitrogen) for 24 h. Following transfection, HEK293 cells were treated with 50 μM MG132 in 10 mL 10% FBS/DMEM for 6 h. Cells were washed, then scraped in PBS and centrifuged at 2,000 *g* for 5 min. Pellets were lysed in RIPA buffer supplemented with 2.5 mM MgCl_2_, 5 mM EDTA, and complete ULTRA protease inhibitor tablet (Roche, 1 tablet per 10 mL) via sonication for 10 seconds at 20% power, then rested on ice for 30 min. The protein concentration was quantified using Pierce Bicinchoninic Acid Protein Kit (Thermo Fisher Scientific) and normalised across conditions. The normalised protein was loaded onto normal mouse IgG conjugated Protein G Dynabeads (Thermo Fisher Scientific) for preclearing. 5 μg anti-V5 antibody (Invitrogen, R960) was conjugated to Protein G Dynabeads (Thermo Fisher Scientific) per sample, then protein lysate from the preclearing was loaded and immunoprecipitated overnight at 4°C. The beads were then washed three times with RIPA buffer at 4°C. Bound proteins were eluted off the beads via heating at 95°C for 10 min and vortexing in elution buffer (RIPA buffer supplemented with 4% SDS and 1× Laemmli buffer), before being subjected to western blot.

### Co-Immunoprecipitation

2.5×10^6^ HEK293 cells were seeded in a T-25 flask and transfected the following day with a total of 4 μg plasmid (at a ratio of 3 μg pcDNA3.1 ORMDL3–V5/His : 1 μg pcDNA3.1 EV–Myc/His, pcDNA3.1 Derlin-2–Myc/His or pcDNA3.1 Derlin-3–Myc/His) using 16 μL Lipofectamine 3000 reagent and 8 μL P3000 supplemental reagent (Invitrogen) for 24 h. Following transfection, cells were treated with 50 μM MG132 for 2 h. Cells were washed, then scraped in PBS and centrifuged at 2,000 *g* for 5 min. Pellets were lysed in RIPA buffer supplemented with 2.5 μM MgCl_2_, 5 μM EDTA, and complete ULTRA protease inhibitor tablet (Roche, 1 tablet per 10 mL) via sonication for 10 seconds at 20% power, then rested on ice for 30 min. The protein concentration was quantified using Pierce Bicinchoninic Acid Protein Kit (Thermo Fisher Scientific) and normalised across conditions. The normalised protein lysate was loaded onto V5-Trap Magnetic Agarose beads (Chromotek) and immunoprecipitated overnight at 4°C. The beads were then washed three times with IP wash buffer (50 mM NaCl, 10 mM Tris, pH 7.5) at 4°C. Bound proteins were eluted from the beads via heating at 95°C for 10 min and vortexing in elution buffer (RIPA buffer supplemented with 4% SDS and 1× Laemmli buffer), before being subjected to western blot.

### Proteomic Mass Spectrometry

#### Sample preparations

Samples were lyophilized overnight and reconstituted in 200ul of 6M Guanidine -HCl. The samples were then boiled for 10 minutes followed by 5 minutes cooling at room temperature. The boiling and cooling cycle was repeated a total of 3 cycles. The proteins were precipitated with addition of methanol to final volume of 90% followed by vortex and centrifugation at maximum speed on a benchtop microfuge (14000 rpm) for 10 minutes. The soluble fraction was removed by flipping the tube onto an absorbent surface and tapping to remove any liquid. The pellet was suspended in 200ul of 8 M Urea made in 100mM Tris pH 8.0. TCEP was added to final concentration of 10 mM and Chloro-acetamide solution was added to final concentration of 40 mM and vortex for 5 minutes. 3 volumes of 50mM Tris pH 8.0 were added to the sample to reduce the final urea concentration to 2 M. Trypsin was in 1:50 ratio of trypsin and incubated at 37C for 12 hours. The solution was then acidified using TFA (0.5% TFA final concentration) and mixed. The sample was desalted using C18-StageTips (Thermo) as described by the manufacturer protocol. The peptide concentration of sample was measured using BCA. 20 ug of each sample were used in TMT labeling. TMT10 plex (Thermo P/N 90110) labelling was carried out as described by the manufacturer. After the labeling completion the samples were pooled and the peptides were desalted and fractionated on C18-high pH reverse spin columns. Pierce™ High pH Reversed-Phase Peptide Fractionation Kit (Pierce™ High pH Reversed-Phase Peptide Fractionation Kit Catalog number: 84868) was used. Fractionation protocol as described by the manufacturer kit with the exception that 8 fractions where generated.

#### LC-MS-MS

1.5 ug of each High pH Reversed-Phase fraction was analyzed by ultra-high-pressure liquid chromatography (UPLC) coupled with tandem mass spectroscopy (LC-MS/MS) using nano-spray ionization. The FAIMS nano-spray ionization experiments were performed using an Orbitrap fusion Lumos hybrid mass spectrometer (Thermo) interfaced with nano-scale reversed-phase UPLC (Thermo Vanquish Neo nano UPLC System) using a 50 cm uPAC (Thermo part # COL-NANO050G1B). Peptides were eluted from the C18 column into the mass spectrometer using a linear gradient (8–100%) of ACN (Acetonitrile) at a flow rate of 400 μl/min for 140 min. The buffers used to create the ACN gradient were: Buffer A (99.9% H_2_O, 0.1% formic acid) and Buffer B (80% ACN, 0.1% formic acid). Mass spectrometer parameters are as follows; MS1 survey scan using the orbitrap detector (mass range (m/z): 400-1500 (using quadrupole isolation), 60000 resolution setting, spray voltage of 2200 V, Ion transfer tube temperature of 290 C, AGC target of 400000, and maximum injection time of 50 ms) was followed by data dependent scans (top speed for most intense ions, with charge state set to only include +2-5 ions, and 5 second exclusion time, while selecting ions with minimal intensities of 50000 at in which the collision event was carried out in the high energy collision cell (HCD Collision Energy of 38%) and the first quadrupole isolation window was set at 0.8 (m/z). The fragment masses were analyzed in the orbitrap detector (mass range (m/z): automatic scan with first scan at m/z= 100. The resolution was set at 30000 resolution. AGC Target set to 30000, and maximum injection time: 54 ms. Protein identification and quantification was carried out using Peaks Studio X (Bioinformatics solutions Inc.).

### Proteomic Analyses

Abundance values for each protein were generated for two replicates each of WT, Derlin-1 KO, Derlin-2 KO, Derlin-3 KO, and Derlin TKO. These estimated abundance measures were merged across all samples and used as input to the Perseus software package (Tyanova et al., 2016) for processing. Abundance values were log-transformed and proteins that were not detected in all samples were removed. Then, the values were normalized using a “width adjustment” before being exported to the R statistical environment (www.cran.org). Within R, the data were used in differential abundance analysis using the limma package (Ritchie et al., 2015) to compare the wild-type and each knockout group. Significantly differentially abundant proteins (adjusted p-value <0.05) were used as input for enrichment analysis in the Gene Ontology, KEGG, and Reactome ontologies with the gProfiler tool (https://biit.cs.ut.ee/gprofiler/gost) (Peterson et al., 2020). Multi-dimensional scaling was performed and plotted with functions within limma using all proteins in the dataset and scatter plots (i.e. volcano plots) were made with the ggplot2 package in R.

### Western blotting

Once cells were finished with transfection and/or treatments cells were washed in PBS, scrapped off the surface offthe plate in PBS and centrifuged at 2,000g for 5 min. The supernatant was discarded, and the cell pellet was resuspended in RIPA buffer (Thermo Scientific) supplemented with 2.5 mM MgCl_2_, 5 mM EDTA, and 10% (v/v) protease inhibitor (Sigma), sonicated for 5 seconds at 20% power, then incubated on ice for 30 min. Protein concentration was determined via Pierce Bicinchoninic Acid Protein Kit and normalized in RIPA buffer with 1 × Laemmli buffer. Equal amounts of protein were loaded onto a 6%, 8%, 10%, 12%, 14%, 16%, or 18% (w/v) SDS-PAGE and transferred to a nitrocellulose membrane (Biorad). Total protein was stained for using REVERT 700 (LICOR) according to the manufacturer’s instructions. Membranes were blocked in 5% (w/v) BSA in TBST and probed with the following antibodies in 5% BSA in TBST overnight at 4°C: BiP (Thermo, MA5-27686), PERK (Cell Signaling Technology, 5683), IRE1α (Cell Signaling Technology, 3294), ATF6 (Proteintech, 66563-1-IG), MIC25 (Proteintech, 66597-1-IG), MIC60 (Abcam, Ab110329), Fis1 (Santa Cruz, SC-376447), Drp1 (Cell Signaling Technology, 53945), Mfn2 (Thermo, MA5-27647), ORMDL (Abclonal, A14951), Dj-1 (Cell Signaling Technology, 5933), Grp75 (Enzo, ADI-SPS-825-F), VDAC (Santa Cruz, SC-390996), IP3R1 (Invitrogen, PA1-901), IP3R2 (Santa Cruz, SC-398434), IP3R3 (Thermo, A302-159A), V5 (Invitrogen, R960), Myc (Proteintech, 16286-1-AP), HA (Cell Signaling Technology, 3724), Derlin-1 (Sigma, SAB4200148), and Derlin-2 (Sigma, D1194). After incubation, blots were washed 3 times for 10 min in TBST, then incubated for 1 h with the corresponding secondary antibody (BioRad, 12004161, 12004158, 1705046, or 1705046), then washed 3 times for 10 min in TBST before imaging on the ChemiDoc MP Imaging System. Blots requiring chemiluminescent detection used ECL substrate (BioRad, 1705061 or 1705061).

### Subcellular fractionation

Mitochondria, endoplasmic reticulum (ER), and mitochondria-ER contact sites (MERCs) were isolated from HEK293 cells and were grown to confluence in 90-mm dishes were detached with 1× trypsin, pelleted at 600 × g for 5 min at room temperature, and washed twice with PBS. Cell pellets were resuspended in isolation buffer containing sucrose, mannitol, and EGTA and homogenized. Nuclei and unbroken cells were removed by two sequential centrifugations at 600 × g for 10 min at room temperature. Crude mitochondria were collected by centrifugation at 10,000 × g for 10 min at 4 °C. The resulting supernatant was further centrifuged at 100,000 × g for 1 h at 4 °C to isolate the ER fraction. Crude mitochondria were resuspended in mitochondrial resuspension buffer and layered onto a 30% Percoll gradient. Mitochondria and MERCs fractions were separated by ultracentrifugation at 95,000 × g for 30 min at 4 °C. The mitochondrial fraction, collected as a pellet at the bottom of the gradient, was washed and re-pelleted at 6,300 × g for 10 min at 4 °C. The MERCs fraction, appearing as a diffuse band above the mitochondria, was collected and pelleted at 10,000 × g for 1 h at 4 °C. Fraction purity was verified by immunoblotting using antibodies against TOMM20 (mitochondria; Santa Cruz Biotech. F-10 sc-17764), Sec61B (ER; Proteintech 51020-2-AP), and SIGMA1R (MERCS; Santa Cruz Biotech. B-5 137075). Tagged ORMDLs and Derlins were immunoblotted using antibodies against V5 (Santa Cruz Biotech. C-9 sc-271944) and Myc (Santa Cruz Biotech. 9E10 sc-40).

### Proximity ligation assays

Proximity ligation assays (PLA) were conducted to examine the proximity and potential interaction of two proteins (within < 40 nm). Duolink® In Situ Red Starter Kit (Sigma-Aldrich, # DUO92102) was employed according to the manufacturer’s guidelines. Cells cultured on glass slides underwent fixation, permeabilization, and blocking before overnight incubation with paired primary antibodies (mouse anti-V5) and rabbit anti-VDAC1 (Proteintech, #5259-1-AP). Following a buffer wash, paired secondary antibodies (anti-rabbit PLUS and anti-mouse MINUS) conjugated with oligonucleotides were applied. When the distance between the two proteins of interest was less than 40 nm, ligase action connected the oligonucleotides, forming a closed circular DNA. Signal amplification occurred through rolling circle amplification (RCA) under polymerase action, resulting in dot-like PLA signals observable under a fluorescence microscope. Simultaneous negative control PLA experiments were performed, and PLA signals were quantified using the “Particle Analysis” function of ImageJ.

### Measurement of OCR Using Seahorse

Oxygen consumption rate was measured for WT, Derlin-1 KO, Derlin-2 KO, Derlin-3 KO, Derlin TKO and ORMDL KD HEK293 cells using an XF24 bioanalyzer (Seahorse Bioscience: North Billerica, MA), as previously described (Dranka et al., 2011; Pereira et al., 2017). Cells were seeded at 30,000 per/well. The media was changed to XF-DMEM (supplemented with 1 g/L d-glucose, 0.11 g/L sodium pyruvate, and 4 mM l-glutamine), and cells were incubated without CO^2^ for 60 min. Cells were treated with oligomycin (1 μg/mL), carbonyl cyanide 4-(trifluoromethoxy)phenylhydrazone (FCCP; 1 μM), rotenone (1 μM), and antimycin A (1 μM), in that order, while remaining in the XF-DMEM media. After measurement, cells were lysed accordingly to prior protocols (Pereira et al., 2017) using 20 μL of 10 mM Tris with 0.1% Triton X-100 added at pH 7.4, and media replaced with 480 μL of Bradford reagent. Total protein concentration was then measured at absorbance at 595 nm and used for normalization.

### Measurement of mitochondrial Ca^2+^ uptake

Mitochondrial Ca2+ uptake and efflux were measured using a dual-wavelength emission fluorimeter, following a modified version of the protocol described earlier(Shukla et al., 2025; Tomar et al., 2016). An equal number of cells (2.5 × 10^6^) were first washed with Ca^2+^/Mg^2+^-free DPBS (GIBCO) and then permeabilised in Ca^2+^ free intracellular medium (ICM: 120 mM KCl, 10 mM NaCl, 1 mM KH2PO4, 20 mM HEPES-Tris, pH 7.2) containing 2.5 mM succinate to energize the mitochondria, 20 μg/ml digitonin to permeabilize the plasma membrane, and 1.5 μM thapsigargin to inhibit the SERCA pump. Bath Ca^2+^ concentration was monitored by adding Fura-FF. Fluorescence was monitored using a Delta RAM multi-wavelength excitation, dual-wavelength emission fluorimeter (Delta RAM, PTI), based on the Fura-FF Ca^2+^ bound/unbound fluorescence ratio at 340- and 380-nm excitation/510-nm emission wavelengths. The cell suspension was maintained with continuous stirring at 37°C. At selected intervals, 5 μM Ca^2+^, 1 μM Ru360 (mitochondrial calcium blocker), and 10 μM FCCP (a mitochondrial uncoupler) were added to the suspension.

### Measurement of ER and mitochondrial Ca^2+^ release

To assess calcium release from the endoplasmic reticulum (ER) and mitochondria, cells were prepared as outlined in the “Measurement of mitochondrial Ca^2+^ uptake” section and resuspended in Ca^2+^ free ICM buffer containing 2.5 mM succinate, 20 μg/ml digitonin, 1 μM Ru360 (a mitochondrial calcium blocker), and 10 μM CGP (a selective inhibitor of the mitochondrial sodium-calcium exchanger). Bath Ca^2+^ concentration was monitored by the ratiometric Ca^2+^ sensor Fura-FF using a Delta RAM multi-wavelength excitation, dual-wavelength emission fluorimeter (Delta RAM, PTI). After baseline recordings, 2 μM thapsigargin was added at the indicated time point to inhibit SERCA to measure the ER calcium release. Subsequently, 10 μM FCCP was added to dissipate the mitochondrial membrane potential (Δψm) and measure the release of mitochondrial matrix free Ca^2+^.

### Measurement of _m_Ca^2+^ retention capacity (CRC)

To measure mitochondrial Ca^2+^ retention capacity (CRC), 2.5 × 10^6^ cells prepared as described in the “Measurement of mitochondrial Ca^2+^ uptake” section and resuspended in a Ca^2+^ free ICM buffer containing 2.5 mM succinate, 20 μg/ml digitonin and 1.5 μM thapsigargin. The permeabilised cells were incubated with the fluorometric calcium indicator Fura-FF. Fluorescence was recorded using a spectrofluorometer (Delta RAM, Photon Technology International). After baseline recordings, sequential Ca^2+^ boluses (5 μM each) were added at the indicated time points. Once mitochondrial Ca^2+^ uptake is completely ablated and a steady state was observed, 10 μM FCCP was added to dissipate the Δψm and release matrix-free Ca^2+^. The number of Ca^2+^ pulses taken up by the cells was counted to determine mitochondrial CRC.

### Lipidomics Profiling

#### Cell extraction for lipids

10×10^7^ cells per cell line were mixed with 1 mL of Extraction Buffer containing IPA/H2O/Ethyl Acetate (30:10:60, v/v/v) and Avanti Lipidomix Internal Standard (diluted 1:1000) (Avanti Polar Lipids, Inc. Alabaster, AL). Samples were vortexed and transferred to bead mill tubes for homogenization using a VWR Bead Mill at 6000 g for 30 seconds, repeated twice. The samples were then sonicated for 5 minutes and centrifuged at 15,000 g for 5 minutes at 4°C. The upper phase was transferred to a new tube and kept at 4°C. To re-extract the tissues, another 1 mL of Extraction Buffer (30:10:60, v/v/v) was added to the plasma pellet-containing tube. The samples were vortexed, homogenized, sonicated, and centrifuged as described earlier. The supernatants from both extractions were combined, and the organic phase was dried under liquid nitrogen gas.

#### Sample reconstitution for lipids

The dried samples were reconstituted in 300 µL of Solvent A (IPA/ACN/H2O, 45:35:20, v/v/v). After brief vortexing, the samples were sonicated for 7 minutes and centrifuged at 15,000 g for 10 minutes at 4°C. The supernatants were transferred to clean tubes and centrifuged again for 5 minutes at 15,000 g at 4°C to remove any remaining particulates. For LC-MS lipidomic analysis, 60 µL of the sample extracts were transferred to mass spectrometry vials.

#### LC-MS analysis for lipids

Sample analysis was performed within 36 hours after extraction using a Vanquish UHPLC system coupled with an Orbitrap Exploris 240™ mass spectrometer equipped with a H-ESI™ ion source (all Thermo Fisher Scientific). A Waters (Milford, MA) CSH C18 column (1.0 × 150 mm × 1.7 µm particle size) was used. Solvent A consisted of ACN:H2O (60:40; v/v) with 10 mM Ammonium formate and 0.1% formic acid, while solvent B contained IPA:ACN (95:5; v/v) with 10 mM Ammonium formate and 0.1% formic acid. The mobile phase flow rate was set at 0.11 mL/min, and the column temperature was maintained at 65 °C. The gradient for solvent B was as follows: 0 min 15% (B), 0–2 min 30% (B), 2–2.5 min 48% (B), 2.5–11 min 82%, 11–11.01 min 99% (B), 11.01–12.95 min 99% (B), 12.95–13 min 15% (B), and 13–15 min 15% (B). Ion source spray voltages were set at 4,000 V and 3,000 V in positive and negative mode, respectively. Full scan mass spectrometry was conducted with a scan range from 200 to 1000 m/z, and AcquireX mode was utilized with a stepped collision energy of 30% with a 5% spread for fragment ion MS/MS scan.

### Metabolomics Profiling

400,000 cells per cell line were extracted in -80°C 80:20 methanol:water, vortexed, incubated on dry ice for 10 minutes, and centrifuged at 16,000 g for 25 minutes, with the supernatant used for LC-MS analysis. Extracts were analyzed within 24 hours by liquid chromatography coupled to a mass spectrometer (LC-MS). The LC–MS method was based on hydrophilic interaction chromatography (HILIC) coupled to the Orbitrap Exploris 240 mass spectrometer (Thermo Scientific) (Wang et al., 2019). The LC separation was performed on a XBridge BEH Amide column (2.1 x 150 mm, 3.5 𝜇m particle size, Waters, Milford, MA). Solvent A is 95%: 5% H2O: acetonitrile with 20 mM ammonium acetate and 20mM ammonium hydroxide, and solvent B is 90%: 10% acetonitrile: H2O with 20 mM ammonium acetate and 20mM ammonium hydroxide. The gradient was 0 min, 90% B; 2 min, 90% B; 3 min, 75% B; 5 min, 75% B; 6 min, 75% B; 7 min, 75% B; 8 min, 70% B; 9 min, 70% B; 10 min, 50% B; 12 min, 50% B; 13 min, 25% B; 14min, 25% B; 16 min, 0% B; 18 min, 0% B; 20 min, 0% B; 21 min, 90% B; 25 min, 90% B. The following parameters were maintained during the LC analysis: flow rate 150 mL/min, column temperature 25 °C, injection volume 5 µL and autosampler temperature was 5 °C. For the detection of metabolites, the mass spectrometer was operated in both negative and positive ion mode. The following parameters were maintained during the MS analysis: resolution of 180,000 at m/z 200, automatic gain control (AGC) target at 3e6, maximum injection time of 30 ms and scan range of m/z 70-1000. Raw LC/MS data were converted to mzXML format using the command line “msconvert” utility (Adusumilli & Mallick, 2017). Data were analyzed via the EL-MAVEN software version 12.

### Metabolomic Analysis

Core metabolomic data analyses were performed in R (v4.4.3; R Core Team, 2025). Cellular extract and media metabolomics data were analyzed separately but using identical analytical procedures, unless otherwise noted. All analyses were conducted using custom R scripts (see Code Availability). *Processing:* Before any analysis was performed, data were processed and normalized. Briefly, Negative values were treated as missing, and zero or missing values were imputed as one-half of the minimum non-zero value per metabolite; metabolites lacking positive measurements were excluded. Data were log2-transformed, and duplicate features were collapsed by retaining the feature with the highest mean abundance.

#### PCA

For each sample type, principal component analysis (PCA) was performed on the log2-transformed metabolite data using the “prcomp” function in base R, with centering and scaling enabled. Prior to PCA, metabolites with zero variance across samples were removed. PCA scores were visualized using the first two principal components, with samples colored by experimental group.

#### Differential Abundance Analysis

Differential metabolite abundance was assessed using linear models with empirical Bayes moderation, as implemented in the *limma* package(Ritchie et al., 2015) or each dataset, metabolite abundances were modeled as a function of experimental group, with group means estimated explicitly for each condition. This model parameterization facilitated direct and biologically interpretable contrasts between experimental groups. Pairwise contrasts were specified to compare each knockout line (D1, D2, D3, and DT) against WT, as well as an aggregate contrast comparing the mean abundance across all knockout lines to WT. Linear models were fit using “lmFit”, followed by contrast estimation and empirical Bayes moderation using “contrast.fit” and “eBayes”. Multiple testing correction was performed using the Benjamini–Hochberg false discovery rate (FDR) procedure implemented in *limma*, with an FDR < 0.05 used to define statistical significance. Heatmaps were generated using the *pheatmap* (Kolde, 2025) package in R to visualize metabolites showing significant differential abundance. Log2-transformed abundances were row-scaled (Z-scored) prior to visualization, and metabolites with zero variance were excluded. Samples were ordered by experimental group, and hierarchical clustering was performed using Euclidean distance and complete linkage. Heatmaps were generated both across all experimental groups and for individual contrasts, displaying either all groups or only those relevant to a given comparison. Where applicable, metabolites were annotated by the direction of change relative to WT.

#### Enrichment Analysis

Metabolite set enrichment analysis was performed using the MetaboAnalyst Quantitative Enrichment Analysis (QEA) framework (Pang et al., 2021). For each contrast, log2-transformed metabolite abundance matrices and corresponding class labels were exported in the required MetaboAnalyst format. Pathway-level enrichment results were returned with FDR adjusted P-values. Enrichment results were imported back into R for downstream processing and visualization in the form of dot plots.

## Supplemental Figure Legends

**Supplemental Figure 1.**
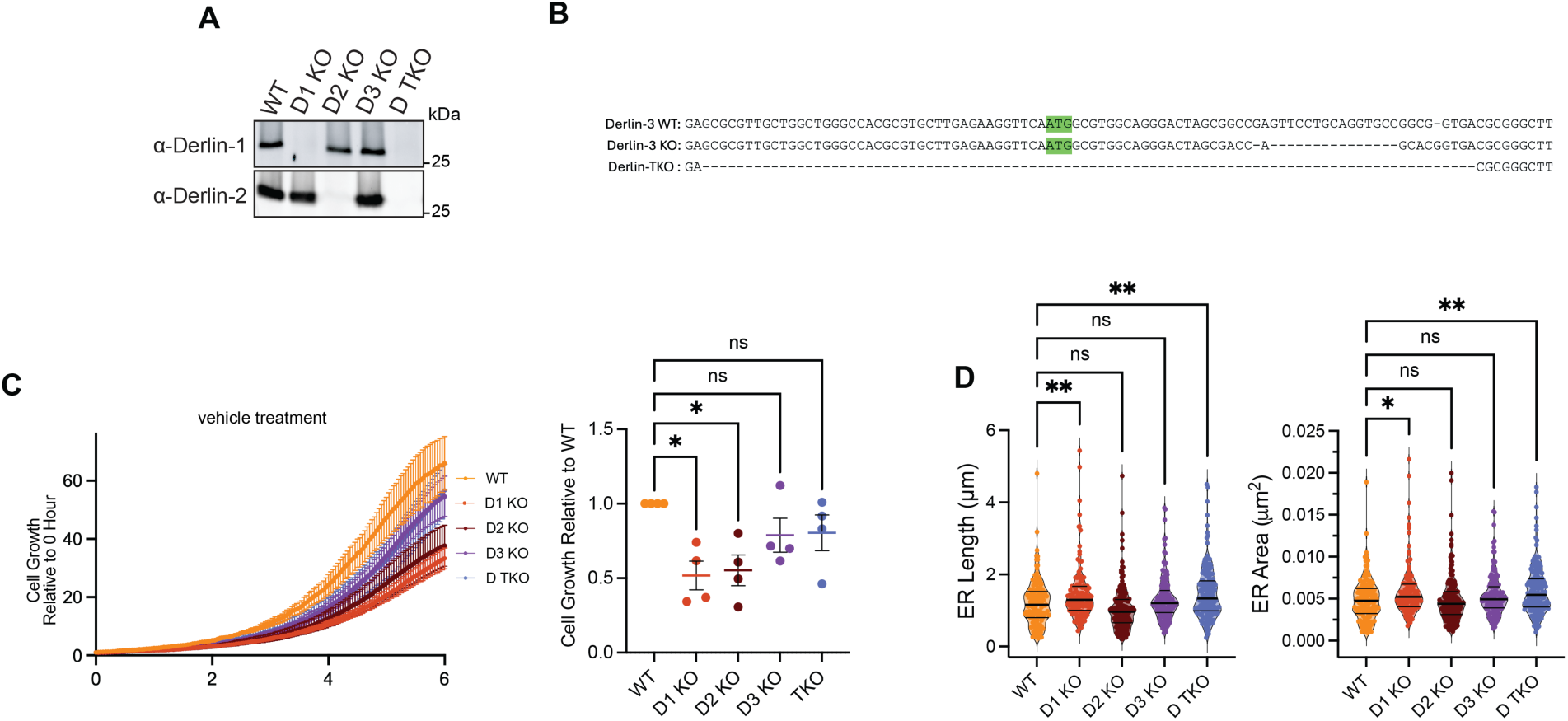
Validation of Derlin KO lines and quantitative analysis of ER and mitochondrial morphology. **(A)** Validation of WT, Derlin-1 KO, Derlin-2 KO, and Derlin TKO HEK293 cells via immunoblotting with endogenous Derlin-1 and Derlin-2 antibodies. Data are representative of *n* = 3 independent experiments. **(B)** Validation of Derlin-3 KO and Derlin TKO HEK293 cells via TOPO cloning of DERLIN-3 genomic locus. **(C)** Proliferation of HEK293 WT and Derlin KO cells treated with vehicle. Confluence was measured as in **Figure 1D**. AUC was quantified and compared to WT. Representative of *n* = 4 independent experiments. Statistical significance was determined by one-way ANOVA followed by Tukey’s multiple-comparison test. ns, not significant. **(D)** Quantitative analyses of ER area and perimeter were performed from TEM images using standardized criteria across all conditions. Each data point represents an individual ER. Statistical significance was determined using one-way ANOVA with multiple-comparison correction. *P < 0.05, **P < 0.01, ***P < 0.001, ****P < 0.0001; ns, not significant.

**Supplemental Figure 2.**
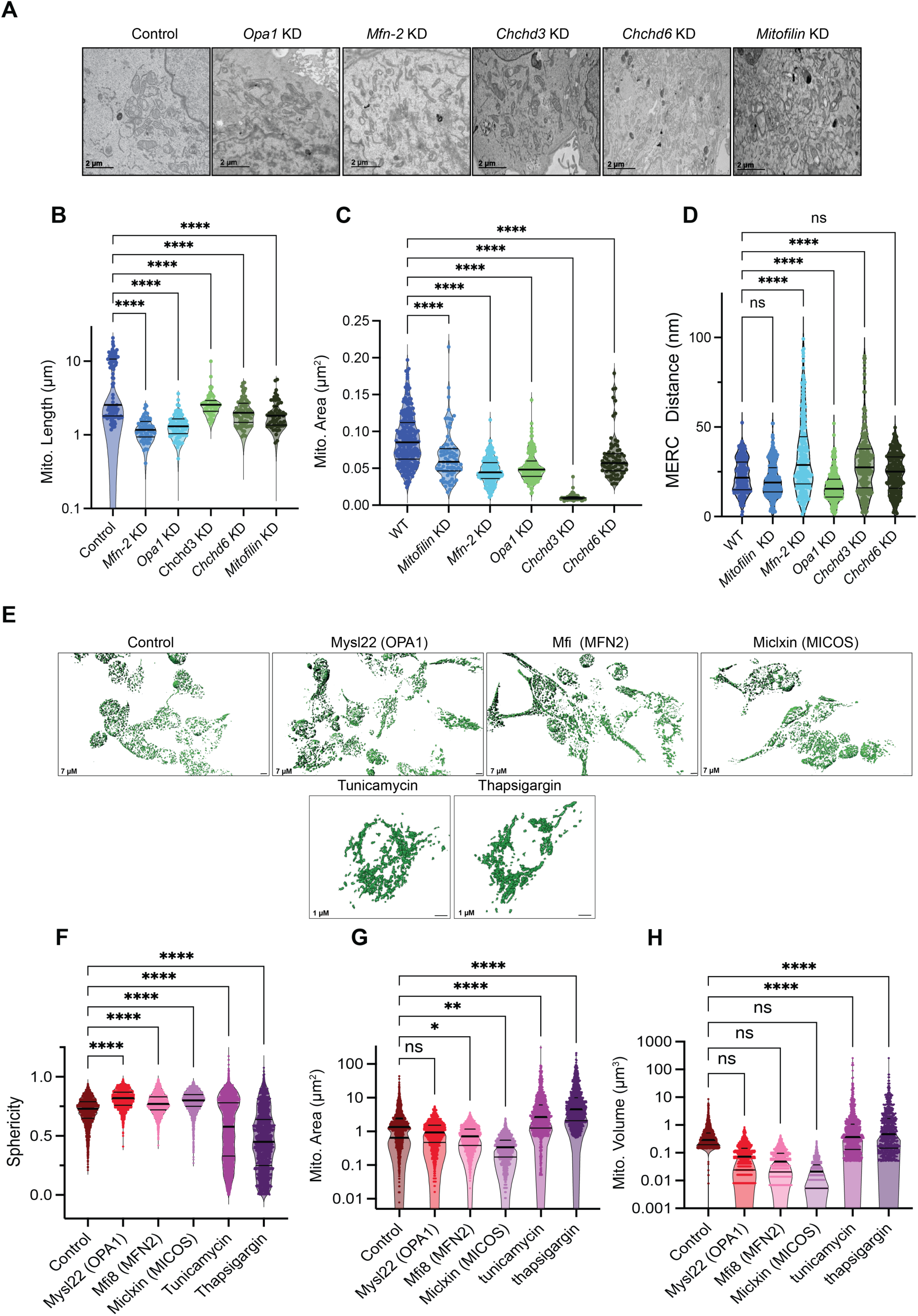
Loss of Derlins does not affect ER and mitochondrial Ca²⁺ release. **(A)** Representative TEM images are shown for Control, *OPA1* knockdown, *MFN2* knockdown, *CHCHD3* knockdown, *CHCHD6* knockdown, and *Mitofilin* knockdown. Mitochondria displays distinct alterations in size and shape depending on the targeted pathway: scale bars, 2 µm. **(B-D)** Quantitative analyses of mitochondrial length (B), mitochondrial area (C), and mitochondria–ER contact (MERC) distance (D) were performed from TEM images using standardized criteria across all conditions from (A). Each data point represents an individual mitochondrion. Statistical significance was determined using one-way ANOVA with multiple-comparison correction. ***P < 0.001, ****P < 0.0001; ns, not significant. **(E)** Representative 3D surface renderings of mitochondria reconstructed from high-resolution confocal z-stack imaging are shown for Control, Mys22, Mfi8, Miclixin, Tunicamycin, and Thapsigargin treatment in HEK293 cells. Mitochondria are displayed as green surface renderings following automated segmentation and batch surface generation, with identical parameters across all treatment groups; scale bar, 7 µm. **(F-H)** Quantification of mitochondrial sphericity (F), mitochondrial area (G), and mitochondrial volume **(H)**. Each dot represents an individual mitochondrion. Inhibitor treatments significantly altered mitochondrial size and shape distributions compared with vehicle control, with all Derlin KOs producing the most pronounced reductions in mitochondrial area and volume and a shift toward increased sphericity, consistent with mitochondrial fragmentation. Statistical significance was determined using one-way ANOVA with appropriate multiple-comparison correction. *P < 0.05, **P < 0.01, ****P < 0.0001; ns, not significant.

**Supplemental Figure 3.**
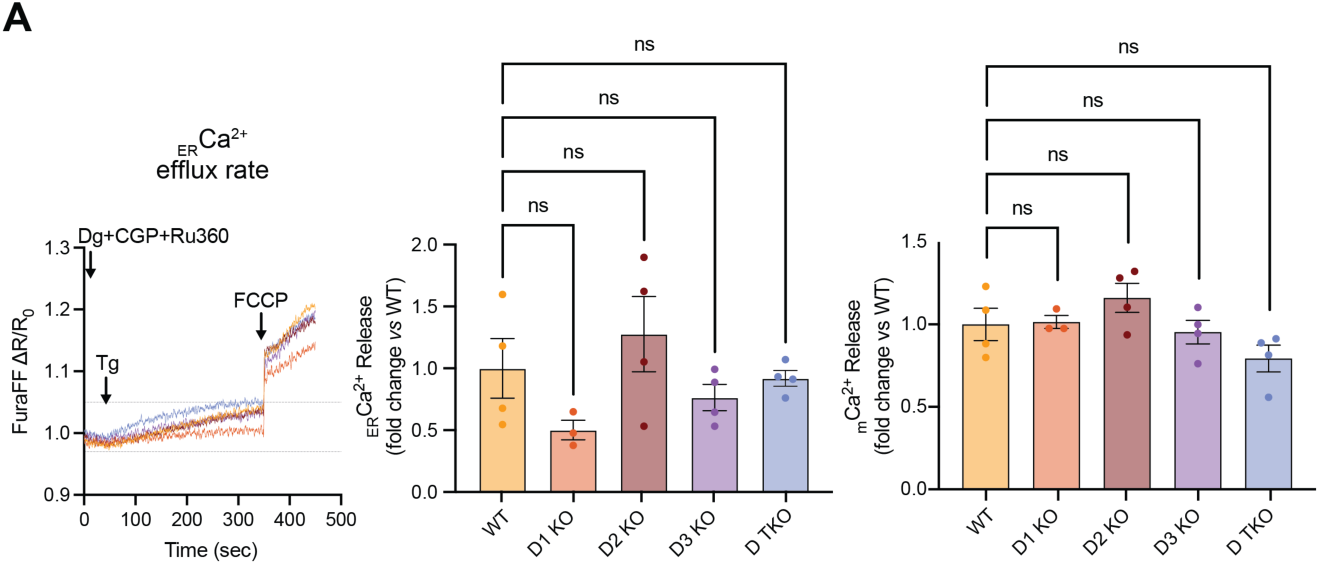
Loss of Derlins does not affect ER and mitochondrial Ca²⁺ release. **(A)** Cells were permeabilized and monitored in Ca²⁺-free buffer using the ratiometric Ca²⁺ indicator Fura-FF. ER Ca²⁺ release was induced by thapsigargin, followed by FCCP treatment to trigger release of mitochondrial matrix Ca²⁺. Data represent *n* = 3–4 independent experiments (mean ± SEM). Statistical significance was determined using one-way ANOVA followed by Dunnett’s multiple-comparison test. ns, not significant.

**Supplemental Figure 4.**
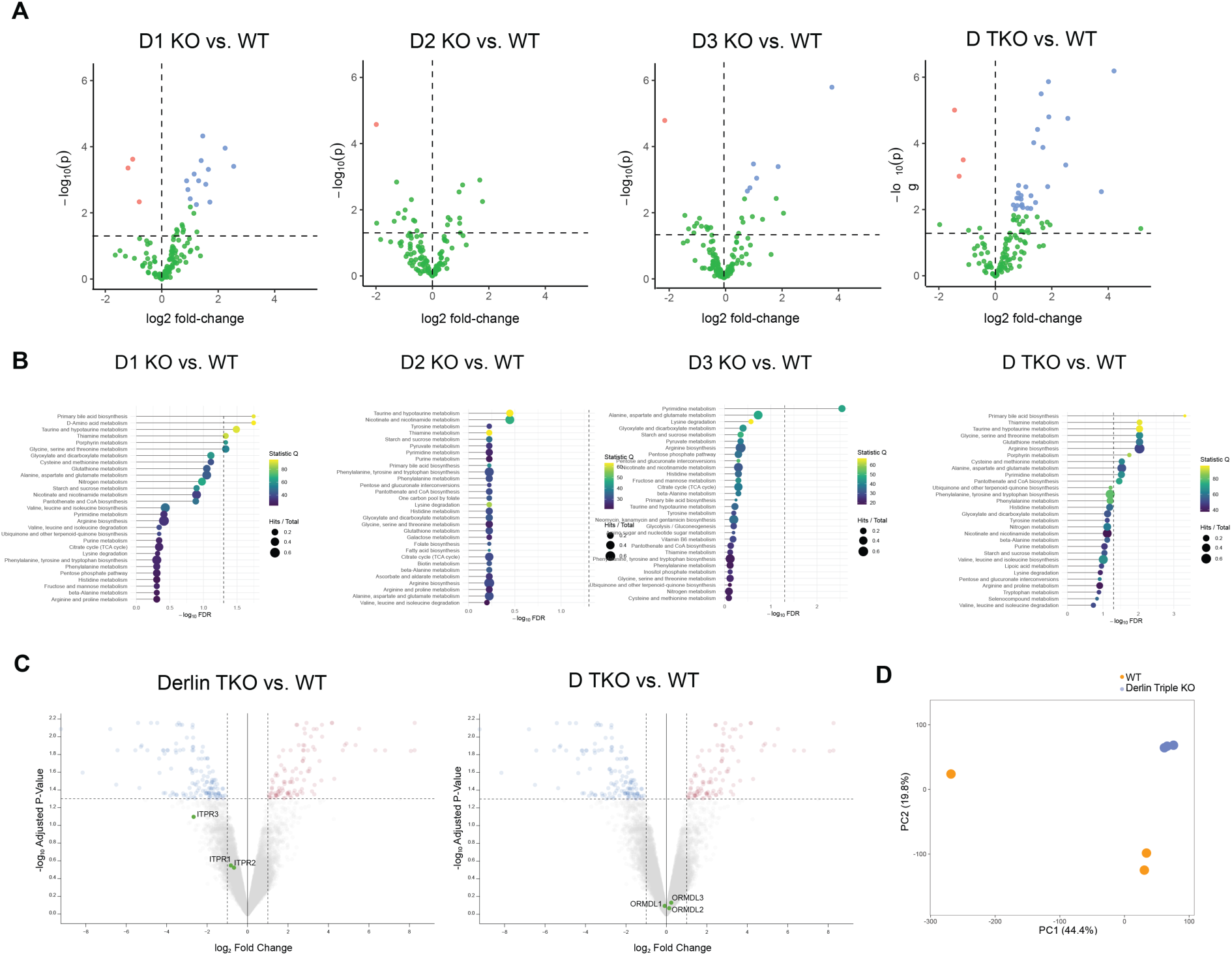
Global metabolomic alterations and pathway enrichment in Derlin KO HEK293 cells. **(A)** Volcano plots showing changes in metabolite abundance between WT and Derlin knockout cells. Metabolites with significantly increased abundance are shown in red, while significantly decreased metabolites are shown in blue. Non-significant metabolites are shown in green. **(B)** Metabolite set enrichment analysis of WT and Derlin knockout cells. Quantitative enrichment analysis (QEA) was performed using MetaboAnalyst on log₂-transformed metabolite abundance data, with pathway enrichment significance reported as FDR-adjusted P values and visualized as dot plots. **(C)** Volcano plots showing changes in RNA abundance between WT and Derlin triple knockout cells. RNA with significantly increased abundance are shown in red, while those with significantly decreased abundance are shown in blue. Non-significant changes in RNA abundance are shown in gray. All ORMDLs and ITPRs are highlighted in green. **(D)** Principal component analysis (PCA) was performed on RNA-Seq count per million reads from WT and Derlin triple KO HEK293 cells. Data were centered and scaled, and samples are shown along the first two principal components, colored by their respective genotypes.

**Supplemental Figure 5.**
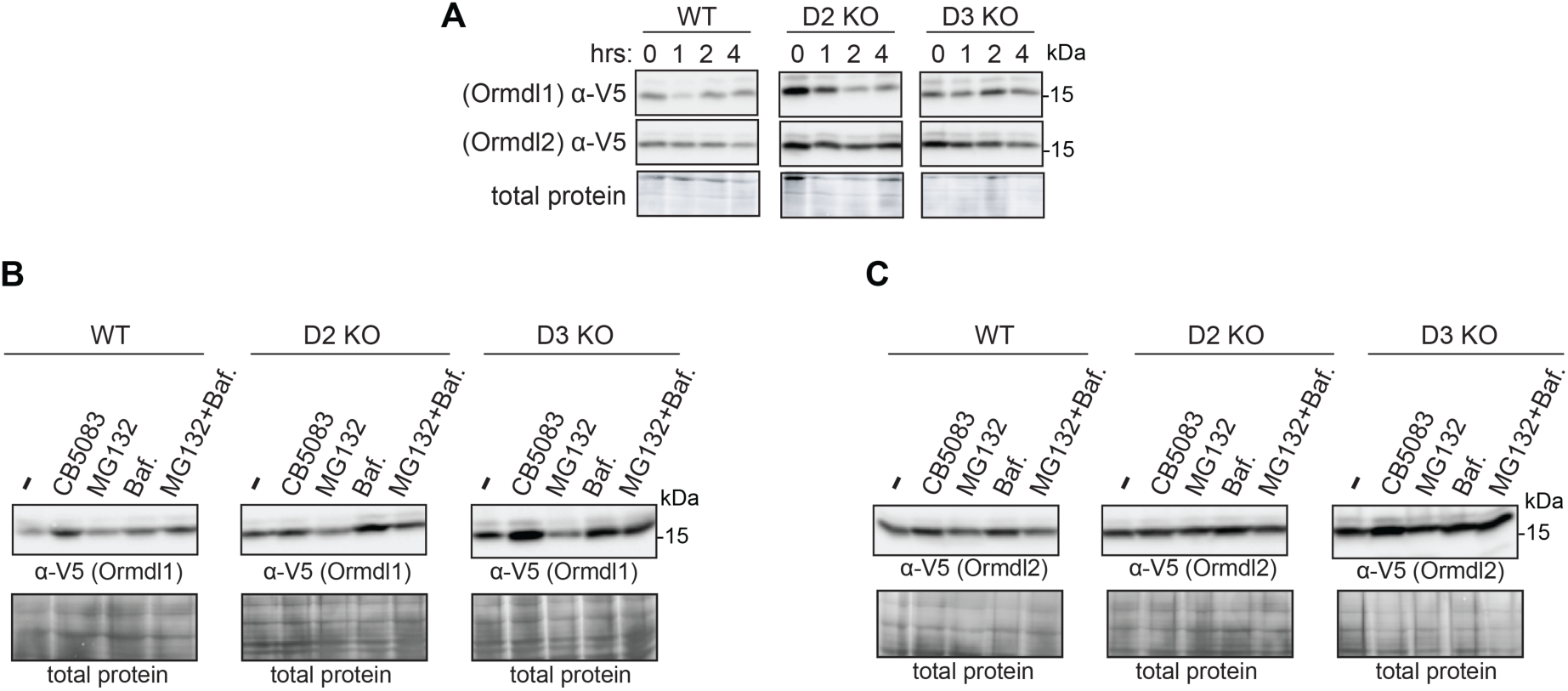
Derlin-2 and Derlin-3 dependent regulation of ORMDL1 and ORMDL2 levels. **(A)** WT, Derlin-2 KO, and Derlin-3 KO HEK293 cells were transfected with ORMDL1-V5/His or ORMDL2-V5/His for 24 h, then treated with cycloheximide (10 µg/mL) for the indicated times up to 4 h. Protein stability was assessed by western blot. Data are representative of *n* = 3 independent experiments. **(B-C)** WT, Derlin-2 KO, and Derlin-3 KO, and Derlin TKO cells were transfected with ORMDL1-V5/His or ORMDL2-V5/His for 24 h, then treated for 6 h with vehicle, the p97/VCP inhibitor CB5083, the proteasome inhibitor MG132, or the lysosomal inhibitor bafilomycin. Protein levels were analyzed by western blot. Data are representative of *n* = 3 independent experiments.

**Supplemental Figure 6.**
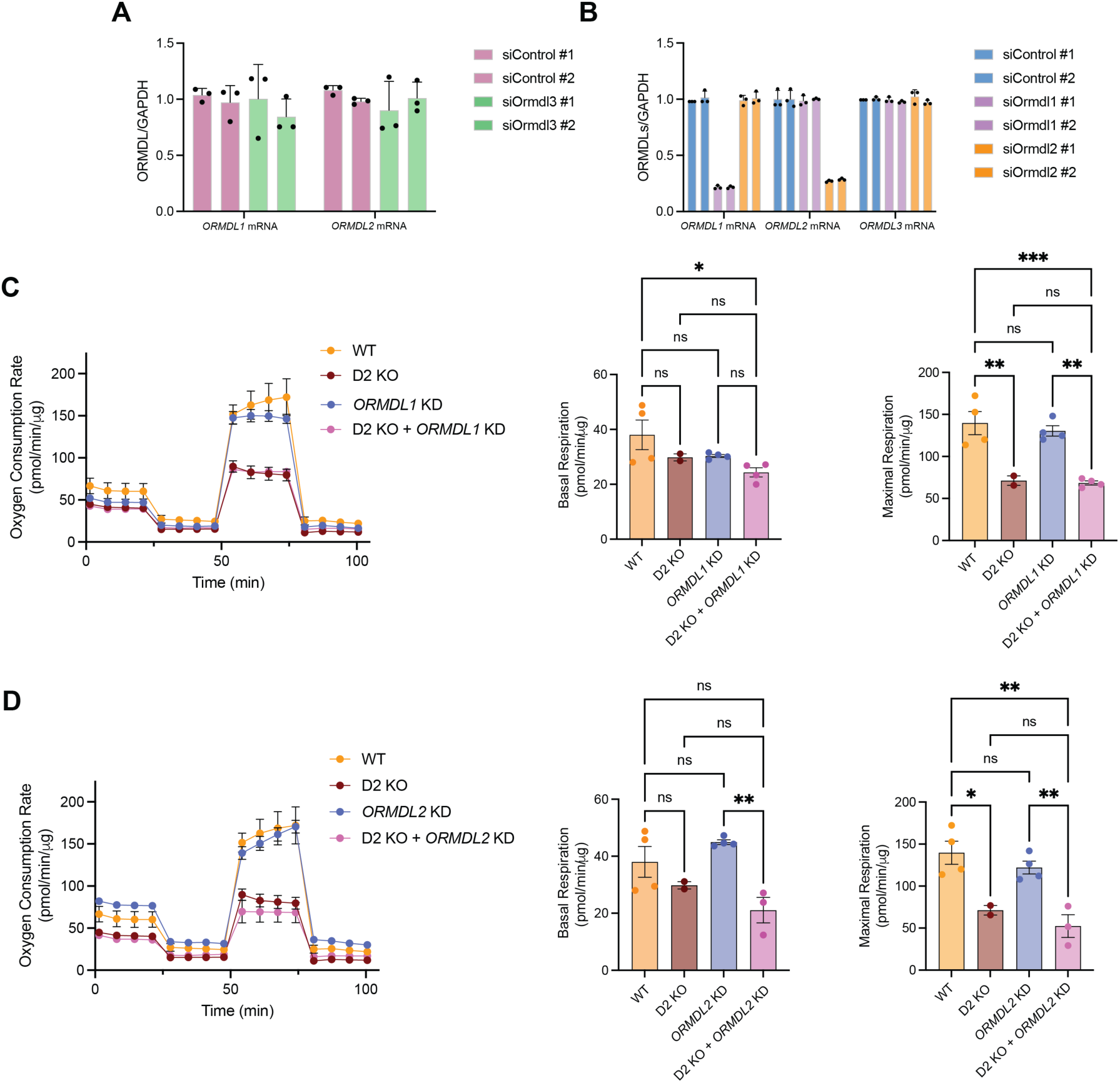
ORMDL1 KD or ORMDL2 KD in Derlin-2 KOs does not rescue mitochondria respiration function. **(A)** ORMDL1 and ORMDL2 mRNA levels were measured by quantitative RT–PCR in cells transfected with two independent control siRNAs (siControl #1 and #2) or two independent ORMDL3-targeting siRNAs (siORMDL3 #1 and #2). ORMDL expression was normalized to GAPDH and is shown relative to control conditions. Data are presented as mean ± SEM from independent experiments, with individual data points representing biological replicates. **(B)** ORMDL1, ORMDL2, and ORMDL3 mRNA levels were measured by quantitative RT–PCR in cells transfected with two independent control siRNAs (siControl #1 and #2) or two independent ORMDL-targeting siRNAs (siORMDL1 or siORMDL2). ORMDL expression was normalized to GAPDH and is shown relative to control conditions. Data are presented as mean ± SEM from independent experiments, with individual data points representing biological replicates.**(C)** Oxygen consumption rate was measured by Seahorse extracellular flux analysis in WT, Derlin-2 KO, ORMDL1 KD, and Derlin-2 KO with ORMDL1 KD HEK293 cells. Basal and maximal respiration were quantified following sequential injection of oligomycin, FCCP, and rotenone/antimycin A. Data are shown as mean ± SEM from independent experiments, with individual data points representing biological replicates. Statistical significance was determined by one-way ANOVA followed by Tukey’s multiple-comparison test. *P < 0.05, **P < 0.01, ***P < 0.001; ns, not significant. **(D)** Oxygen consumption rate was measured by Seahorse extracellular flux analysis in WT, Derlin-2 KO, ORMDL2 KD, and Derlin-2 KO with ORMDL2 KD HEK293 cells. Basal and maximal respiration were quantified following sequential injection of oligomycin, FCCP, and rotenone/antimycin A. Data are shown as mean ± SEM from independent experiments, with individual data points representing biological replicates. Statistical significance was determined by one-way ANOVA followed by Tukey’s multiple-comparison test. *P < 0.05, **P < 0.01, ***P < 0.001.

**Supplemental Figure 7.**
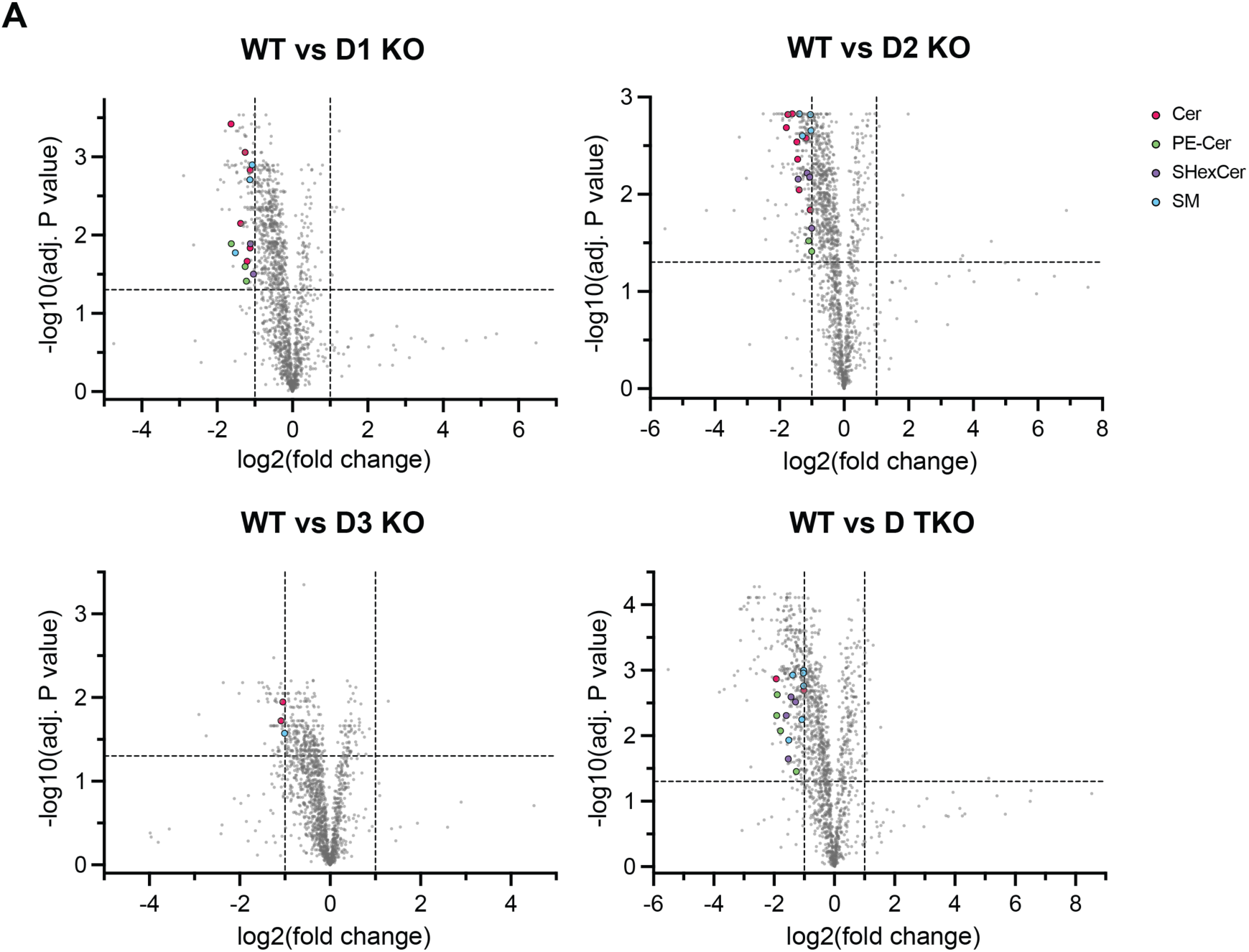
Global lipidomic changes in Derlin-deficient HEK293 cells. **(A)** Volcano plots demonstrating changes to lipids levels versus significance identified in HEK Derlin KO cell lines compared to WT. Sphingolipids that were significantly differentially expressed are colored according to lipid class. Cer, ceramides; PE-Cer, ceramide phosphoethanolamine; SHexCer, sulfated hexoceramide; SM, sphingomyelin. Significantly, different lipid classes have adjusted p-values < 0.05 to correct for multiple comparisons using FDR, with a log fold change less than -1 or greater than 1.

## References

1. Acevedo, N., Reinius, L. E., Greco, D., Gref, A., Orsmark-Pietras, C., Persson, H., Pershagen, G., Hedlin, G., Melén, E., Scheynius, A., Kere, J., & Söderhäll, C. (2014). Risk of childhood asthma is associated with CpG-site polymorphisms, regional DNA methylation and mRNA levels at the GSDMB/ORMDL3 locus. Human Molecular Genetics, 24(3), 875. 10.1093/HMG/DDU479

2. Adams, B. M., Oster, M. E., & Hebert, D. N. (2019). Protein Quality Control in the Endoplasmic Reticulum. The Protein Journal. 10.1007/s10930-019-09831-w

3. Adusumilli, R., & Mallick, P. (2017). Data Conversion with ProteoWizard msConvert. Methods in Molecular Biology (Clifton, N.J.), 1550, 339–368. 10.1007/978-1-4939-6747-6_23

4. Ainbinder, A., Boncompagni, S., Protasi, F., & Dirksen, R. T. (2023). Call to action to properly utilize electron microscopy to measure organelles to monitor disease. European Journal of Cell Biology, 102(4), 151365. 10.1016/j.ceca.2014.11.002

5. An, J., Shi, J., He, Q., Lui, K., Liu, Y., Huang, Y., & Sheikh, M. S. (2012). CHCM1/CHCHD6, Novel Mitochondrial Protein Linked to Regulation of Mitofilin and Mitochondrial Cristae Morphology. The Journal of Biological Chemistry, 287(10), 7411. 10.1074/JBC.M111.277103

6. Bays, N. W., Wilhovsky, S. K., Goradia, a, Hodgkiss-Harlow, K., & Hampton, R. Y. (2001). HRD4/NPL4 is required for the proteasomal processing of ubiquitinated ER proteins. Molecular Biology of the Cell, 12(12), 4114–4128. http://www.pubmedcentral.nih.gov/articlerender.fcgi?artid=60780&tool=pmcentrez&rendertype=abstract

7. Bhaduri, S., Aguayo, A., Ohno, Y., Proietto, M., Jung, J., Wang, I., Kandel, R., Singh, N., Ibrahim, I., Fulzele, A., Bennett, E., Kihara, A., & Neal, S. E. (2022). An ERAD-independent role for rhomboid pseudoprotease Dfm1 in mediating sphingolipid homeostasis. BioRxiv, 2022.07.30.502165. 10.1101/2022.07.30.502165

8. Bhaduri, S., Scott, N. A., & Neal, S. E. (2022). The Role of the Rhomboid Superfamily in ER Protein Quality Control : From Mechanisms and Functions to Diseases. 1–16. 10.1101/cshperspect.a041248

9. Bodnar, N. O., & Rapoport, T. A. (2017). Molecular Mechanism of Substrate Processing by the Cdc48 ATPase Complex. Cell, 169(4), 722–735.e9. 10.1016/j.cell.2017.04.020

10. Brown, R. D. R., & Spiegel, S. (2023). ORMDL in metabolic health and disease. Pharmacology and Therapeutics, 245. 10.1016/j.pharmthera.2023.108401

11. Cipolat, S., De Brito, O. M., Dal Zilio, B., & Scorrano, L. (2004). OPA1 requires mitofusin 1 to promote mitochondrial fusion. Proceedings of the National Academy of Sciences of the United States of America, 101(45), 15927–15932. 10.1073/PNAS.0407043101

12. Davis, D., Kannan, M., & Wattenberg, B. (2018a). Orm/ORMDL proteins: Gate guardians and master regulators. Advances in Biological Regulation, 70(August), 3–18. 10.1016/j.jbior.2018.08.002

13. Davis, D., Kannan, M., & Wattenberg, B. (2018b). Orm/ORMDL proteins: Gate Guardians and Master Regulators. Advances in Biological Regulation, 70, 3. 10.1016/J.JBIOR.2018.08.002

14. Davis, D., Kannan, M., & Wattenberg, B. (2018c). Orm/ORMDL proteins: Gate Guardians and Master Regulators. Advances in Biological Regulation, 70, 3. 10.1016/j.jbior.2018.08.002

15. Davis, D., Kannan, M., & Wattenberg, B. (2018d). Orm/ORMDL proteins: Gate guardians and master regulators. Advances in Biological Regulation, 70, 3–18. 10.1016/j.jbior.2018.08.002

16. Dematteis, G., Tapella, L., Casali, C., Talmon, M., Tonelli, E., Reano, S., Ariotti, A., Pessolano, E., Malecka, J., Chrostek, G., Kulkovienė, G., Umbrasas, D., Distasi, C., Grilli, M., Ladds, G., Filigheddu, N., Fresu, L. G., Mikoshiba, K., Matute, C., … Lim, D. (2024). ER-mitochondria distance is a critical parameter for efficient mitochondrial Ca2+ uptake and oxidative metabolism. Communications Biology 2024 7:1, 7(1), 1294-. 10.1038/s42003-024-06933-9

17. Dougan, S. K., Hu, C.-C. A., Paquet, M.-E., Greenblatt, M. B., Kim, J., Lilley, B. N., Watson, N., & Ploegh, H. L. (2011). Derlin-2-Deficient Mice Reveal an Essential Role for Protein Dislocation in Chondrocytes. Molecular and Cellular Biology, 31(6), 1145–1159. 10.1128/MCB.00967-10

18. Dranka, B. P., Benavides, G. A., Diers, A. R., Giordano, S., Zelickson, B. R., Reily, C., Zou, L., Chatham, J. C., Hill, B. G., Zhang, J., Landar, A., & Darley-Usmar, V. M. (2011). Assessing bioenergetic function in response to oxidative stress by metabolic profiling. Free Radical Biology and Medicine, 51(9), 1621–1635. 10.1016/j.freeradbiomed.2011.08.005

19. Ebert, A. C., Hepowit, N. L., Martinez, T. A., Vollmer, H., Singkhek, H. L., Frazier, K. D., Kantejeva, S. A., Patel, M. R., & MacGurn, J. A. (2025). Sphingolipid metabolism drives mitochondria remodeling during aging and oxidative stress. BioRxiv, 2025.02.26.640157. 10.1101/2025.02.26.640157

20. Eura, Y., Yanamoto, H., Arai, Y., Okuda, T., Miyata, T., & Kokame, K. (2012). Derlin-1 Deficiency Is Embryonic Lethal, Derlin-3 Deficiency Appears Normal, and Herp Deficiency Is Intolerant to Glucose Load and Ischemia in Mice. PLoS ONE, 7(3), e34298. 10.1371/journal.pone.0034298

21. Fugio, L. B., Coeli-Lacchini, F. B., & Leopoldino, A. M. (2020). Sphingolipids and Mitochondrial Dynamic. Cells 2020, Vol. 9, Page 581, 9(3), 581. 10.3390/CELLS9030581

22. Greenblatt, E. J., Olzmann, J. a, & Kopito, R. R. (2011). Derlin-1 is a rhomboid pseudoprotease required for the dislocation of mutant α-1 antitrypsin from the endoplasmic reticulum. Nature Structural & Molecular Biology, 18(10), 1147–1152. 10.1038/nsmb.2111

23. Gupta, S. D., Gable, K., Alexaki, A., Chandris, P., Proia, R. L., Dunn, T. M., & Harmon, J. M. (2015). Expression of the ORMDLS, modulators of serine palmitoyltransferase, is regulated by sphingolipids in mammalian cells. Journal of Biological Chemistry, 290(1), 90–98. 10.1074/jbc.M114.588236

24. Hoelen, H., Zaldumbide, A., Van Leeuwen, W. F., Torfs, E. C. W., Engelse, M. A., Hassan, C., Lebbink, R. J., De Koning, E. J., Resssing, M. E., De Ru, A. H., Van Veelen, P. A., Hoeben, R. C., Roep, B. O., & Wiertz, E. J. H. J. (2015). Proteasomal Degradation of Proinsulin Requires Derlin-2, HRD1 and p97. 10.1371/journal.pone.0128206

25. Huang, C.-H., Hsiao, H.-T., Chu, Y.-R., Ye, Y., & Chen, X. (2013). Derlin2 Protein Facilitates HRD1-mediated Retro-translocation of Sonic Hedgehog at the Endoplasmic Reticulum. Journal of Biological Chemistry, 288(35), 25330–25339. 10.1074/jbc.M113.455212

26. Kadowaki, H., Nagai, A., Maruyama, T., Takami, Y., Satrimafitrah, P., Kato, H., Honda, A., Hatta, T., Natsume, T., Sato, T., Kai, H., Ichijo, H., & Nishitoh, H. (2015). Pre-emptive Quality Control Protects the ER from Protein Overload via the Proximity of ERAD Components and SRP. Cell Reports, 13(5), 944–956. 10.1016/j.celrep.2015.09.047

27. Kadowaki, H., Satrimafitrah, P., Takami, Y., & Nishitoh, H. (2018). Molecular mechanism of ER stress-induced pre-emptive quality control involving association of the translocon, Derlin-1, and HRD1. Scientific Reports, 8(1). 10.1038/s41598-018-25724-x

28. Knop, M., Finger, A., Braun, T., Hellmuth, K., & Wolf, D. H. (1996). Der1, a novel protein specifically required for endoplasmic reticulum degradation in yeast. The EMBO Journal, 15(4), 753–763. http://www.pubmedcentral.nih.gov/articlerender.fcgi?artid=450274&tool=pmcentrez&rendertype=abstract

29. Kolde, R. (2025). Pretty Heatmaps [R package pheatmap version 1.0.13]. CRAN: Contributed Packages. 10.32614/CRAN.PACKAGE.PHEATMAP

30. Lam, J., Katti, P., Biete, M., Mungai, M., Ashshareef, S., Neikirk, K., Lopez, E. G., Vue, Z., Christensen, T. A., Beasley, H. K., Rodman, T. A., Murray, S. A., Salisbury, J. L., Glancy, B., Shao, J., Pereira, R. O., Abel, E. D., & Hintonjr., A. (2021). A universal approach to analyzing transmission electron microscopy with imagej. Cells, 10(9), 2177. 10.3390/cells10092177

31. Lilley, B. N., & Ploegh, H. L. (2004a). A membrane protein required for dislocation of misfolded proteins from the ER. Nature, 429(6994), 834–840. 10.1038/nature02592

32. Lilley, B. N., & Ploegh, H. L. (2004b). A membrane protein required for dislocation of misfolded proteins from the ER. Nature, 429(6994), 834–840. 10.1038/nature02592

33. Lin, L., Chen, L., Lin, G., Chen, X., Huang, L., Yang, J., Chen, S., Lin, R., Yang, D., He, F., Qian, D., Zeng, Y., & Xu, Y. (2025). Derlin-3 manipulates the endoplasmic reticulum stress and IgG4 secretion of plasma cells in lung adenocarcinoma. Oncogene, 44(30), 2620–2633. 10.1038/S41388-025-03435-8

34. Ma, X., Qiu, R., Dang, J., Li, J., Hu, Q., Shan, S., Xin, Q., Pan, W., Bian, X., Yuan, Q., Long, F., Liu, N., Li, Y., Gao, F., Zou, C., Gong, Y., & Liu, Q. (2015). ORMDL3 contributes to the risk of atherosclerosis in Chinese Han population and mediates oxidized low-density lipoprotein-induced autophagy in endothelial cells. Scientific Reports, 5. 10.1038/SREP17194

35. Mahawar, U., Davis, D. L., Kannan, M., Suemitsu, J., Oltorik, C. D., Farooq, F., Fulani, R., Weintraub, C., Allegood, J., & Wattenberg, B. (n.d.). The individual isoforms of ORMDL, the regulatory subunit of serine palmitoyltransferase, have distinctive sensitivities to ceramide. 10.1101/2025.03.20.643044

36. McGovern, D. P. B., Gardet, A., Törkvist, L., Goyette, P., Essers, J., Taylor, K. D., Neale, B. M., Ong, R. T. H., Lagacé, C., Li, C., Green, T., Stevens, C. R., Beauchamp, C., Fleshner, P. R., Carlson, M., D’Amato, M., Halfvarson, J., Hibberd, M. L., Lördal, M., … Seielstad, M. (2010). Genome-wide association identifies multiple ulcerative colitis susceptibility loci. Nature Genetics, 42(4), 332–337. 10.1038/NG.549

37. Neal, S., Jaeger, P. A., Duttke, S. H., Benner, C. K., Glass, C., Ideker, T., & Hampton, R. (2018). The Dfm1 Derlin Is Required for ERAD Retrotranslocation of Integral Membrane Proteins. Molecular Cell, 69(2). 10.1016/j.molcel.2017.12.012

38. Neal, S., Mak, R., Bennett, E. J., & Hampton, R. (2017). A Cdc48 “retrochaperone” function is required for the solubility of retrotranslocated, integral membrane Endoplasmic Reticulum-associated Degradation (ERAD-M) substrates. Journal of Biological Chemistry, 292(8). 10.1074/jbc.M116.770610

39. Needham, P. G., & Brodsky, J. L. (2013). How early studies on secreted and membrane protein quality control gave rise to the ER associated degradation (ERAD) pathway: the early history of ERAD. Biochimica et Biophysica Acta, 1833(11), 2447–2457. 10.1016/j.bbamcr.2013.03.018

40. Neikirk, K., Vue, Z., Katti, P., Rodriguez, B. I., Omer, S., Shao, J., Christensen, T., Garza Lopez, E., Marshall, A., Palavicino-Maggio, C. B., Ponce, J., Alghanem, A. F., Vang, L., Barongan, T., Beasley, H. K., Rodman, T., Stephens, D., Mungai, M., Correia, M., … Hinton, A. O. (2023). Systematic Transmission Electron Microscopy-Based Identification and 3D Reconstruction of Cellular Degradation Machinery. Advanced Biology, 7(6). 10.1002/adbi.202200221

41. Niu, C., Cui, H., Sun, X., Yang, Y., Jiang, X., Cong, Z., Zhang, Y., Niu, Z., & He, W. (2025). DERL3 facilitates the progression of clear cell renal cell carcinoma by promoting epithelial-mesenchymal transition via regulation of the TGFB1 pathway. PloS One, 20(4). 10.1371/JOURNAL.PONE.0322172

42. Oda, Y., Okada, T., Yoshida, H., Kaufman, R. J., Nagata, K., & Mori, K. (2006). Derlin-2 and Derlin-3 are regulated by the mammalian unfolded protein response and are required for ER-associated degradation. Journal of Cell Biology, 172(3), 383–393. 10.1083/JCB.200507057

43. Pang, Z., Chong, J., Zhou, G., De Lima Morais, D. A., Chang, L., Barrette, M., Gauthier, C., Jacques, P. É., Li, S., & Xia, J. (2021). MetaboAnalyst 5.0: narrowing the gap between raw spectra and functional insights. Nucleic Acids Research, 49(W1), W388–W396. 10.1093/NAR/GKAB382

44. Pereira, R. O., Tadinada, S. M., Zasadny, F. M., Oliveira, K. J., Pires, K. M. P., Olvera, A., Jeffers, J., Souvenir, R., Mcglauflin, R., Seei, A., Funari, T., Sesaki, H., Potthoff, M. J., Adams, C. M., Anderson, E. J., & Abel, E. D. (2017). OPA1 deficiency promotes secretion of FGF21 from muscle that prevents obesity and insulin resistance. The EMBO Journal, 36(14), 2126–2145. 10.15252/EMBJ.201696179

45. Peterson, H., Kolberg, L., Raudvere, U., Kuzmin, I., & Vilo, J. (2020). gprofiler2 -- an R package for gene list functional enrichment analysis and namespace conversion toolset g: Profiler. F1000Research, 9. 10.12688/F1000RESEARCH.24956.2/DOI

46. Planas-Serra, L., Launay, N., Goicoechea, L., Heron, B., Jou, C., Juliá-Palacios, N., Ruiz, M., Fourcade, S., Casasnovas, C., De La Torre, C., Gelot, A., Marsal, M., Loza-Alvarez, P., García-Cazorla, À., Fatemi, A., Ferrer, I., Portero-Otin, M., Area-Gómez, E., & Pujol, A. (2023). Sphingolipid desaturase DEGS1 is essential for mitochondria-associated membrane integrity. The Journal of Clinical Investigation, 133(10), e162957. 10.1172/JCI162957

47. Qiu, R., Zhang, H., Zhao, H., Li, J., Guo, C., Gong, Y., & Liu, Q. (2013). Genetic variants on 17q21 are associated with ankylosing spondylitis susceptibility and severity in a Chinese Han population. Scandinavian Journal of Rheumatology, 42(6), 469–472. 10.3109/03009742.2013.786755

48. Ren, G., Tardi, N. J., Matsuda, F., Koh, K. H., Ruiz, P., Wei, C., Altintas, M. M., Ploegh, H., & Reiser, J. (2018). Podocytes exhibit a specialized protein quality control employing derlin-2 in kidney disease. American Journal of Physiology - Renal Physiology, 314(3), F471–F482. 10.1152/ajprenal.00691.2016

49. Ritchie, M. E., Phipson, B., Wu, D., Hu, Y., Law, C. W., Shi, W., & Smyth, G. K. (2015). limma powers differential expression analyses for RNA-sequencing and microarray studies. Nucleic Acids Research, 43(7), e47–e47. 10.1093/NAR/GKV007

50. Sato, B. K., & Hampton, R. Y. (2006). Yeast Derlin Dfm I interacts with Cdc48 and functions in ER homeostasis. Yeast, 23(14–15), 1053–1064. 10.1002/yea.1407

51. Sharma, J., Khan, S., Singh, N. C., Sahu, S., Raj, D., Prakash, S., Bandyopadhyay, P., Sarkar, K., Bhosale, V., Chandra, T., Kumaravelu, J., Barthwal, M. K., Gupta, S. K., Srivastava, M., Guha, R., Ammanathan, V., Ghoshal, U. C., Mitra, K., & Lahiri, A. (2024). ORMDL3 regulates NLRP3 inflammasome activation by maintaining ER-mitochondria contacts in human macrophages and dictates ulcerative colitis patient outcome. The Journal of Biological Chemistry, 300(4), 107120. 10.1016/J.JBC.2024.107120

52. Shukla, S., Kadam, A. A., Goyani, S., Jaiswal, N., Samantaray, K., Kashyap, S., Tomar, D., & Jadiya, P. (2025). Cell-type-specific dysregulation of mitochondrial calcium signaling in Alzheimer’s disease. Cell Communication and Signaling : CCS, 23(1). 10.1186/S12964-025-02460-0

53. Smith, M. E., Tippetts, T. S., Brassfield, E. S., Tucker, B. J., Ockey, A., Swensen, A. C., Anthonymuthu, T. S., Washburn, T. D., Kane, D. A., Prince, J. T., & Bikman, B. T. (2013). Mitochondrial fission mediates ceramide-induced metabolic disruption in skeletal muscle. Biochemical Journal, 456(3), 427–439. 10.1042/BJ20130807

54. Su, M., Qiu, F., Li, Y., Che, T., Li, N., & Zhang, S. (2024). Mechanisms of the NAD+ salvage pathway in enhancing skeletal muscle function. Frontiers in Cell and Developmental Biology, 12, 1464815. 10.3389/FCELL.2024.1464815/FULL

55. Sugiyama, T., Murao, N., Kadowaki, H., Takao, K., Miyakawa, T., Matsushita, Y., Katagiri, T., Futatsugi, A., Shinmyo, Y., Kawasaki, H., Sakai, J., Shiomi, K., Nakazato, M., Takeda, K., Mikoshiba, K., Ploegh, H. L., Ichijo, H., & Nishitoh, H. (2021). ERAD components Derlin-1 and Derlin-2 are essential for postnatal brain development and motor function. IScience, 24(7), 102758. 10.1016/j.isci.2021.102758

56. Tan, X., He, X., Jiang, Z., Wang, X., Ma, L., Liu, L., Wang, X., Fan, Z., & Su, D. (2015). Derlin-1 is overexpressed in human colon cancer and promotes cancer cell proliferation. Molecular and Cellular Biochemistry, 408(1–2), 205–213. 10.1007/s11010-015-2496-x

57. Tomar, D., Dong, Z., Shanmughapriya, S., Koch, D. A., Thomas, T., Hoffman, N. E., Timbalia, S. A., Goldman, S. J., Breves, S. L., Corbally, D. P., Nemani, N., Fairweather, J. P., Cutri, A. R., Zhang, X., Song, J., Jaña, F., Huang, J., Barrero, C., Rabinowitz, J. E., … Madesh, M. (2016). MCUR1 Is a Scaffold Factor for the MCU Complex Function and Promotes Mitochondrial Bioenergetics. Cell Reports, 15(8), 1673–1685. 10.1016/j.celrep.2016.04.050

58. Twomey, E. C., Ji, Z., Wales, T. E., Bodnar, N. O., Ficarro, S. B., Marto, J. A., Engen, J. R., & Rapoport, T. A. (n.d.). Substrate processing by the Cdc48 ATPase complex is initiated by ubiquitin unfolding. 10.1126/science.aax1033

59. Tyanova, S., Temu, T., Sinitcyn, P., Carlson, A., Hein, M. Y., Geiger, T., Mann, M., & Cox, J. (2016). The Perseus computational platform for comprehensive analysis of (prote)omics data. Nature Methods 2016 13:9, 13(9), 731–740. 10.1038/nmeth.3901

60. Volpi, V. G., Ferri, C., Fregno, I., Del Carro, U., Bianchi, F., Scapin, C., Pettinato, E., Solda, T., Feltri, M. L., Molinari, M., Wrabetz, L., & D’Antonio, M. (2019). Schwann cells ER-associated degradation contributes to myelin maintenance in adult nerves and limits demyelination in CMT1B mice. PLOS Genetics, 15(4), e1008069. 10.1371/journal.pgen.1008069

61. Wang, H., Yi, Z., Yan, S., Wang, Y., Zhang, W., Guo, R., Li, Y., Wang, R., Li, H., Li, X., & Song, J. (2025). Advances in ORMDL Research in Malignant Tumors: A Review. OncoTargets and Therapy, 18, 821–832. 10.2147/OTT.S537194

62. Wang, J., Hua, H., Ran, Y., Zhang, H., Liu, W., Yang, Z., & Jiang, Y. (2008). Derlin-1 is overexpressed in human breast carcinoma and protects cancer cells from endoplasmic reticulum stress-induced apoptosis. Breast Cancer Research, 10(1). 10.1186/bcr1849

63. Wang, L., Xing, X., Chen, L., Yang, L., Su, X., Rabitz, H., Lu, W., & Rabinowitz, J. D. (2019). Peak Annotation and Verification Engine for Untargeted LC-MS Metabolomics. Analytical Chemistry, 91(3), 1838–1846. 10.1021/ACS.ANALCHEM.8B03132

64. Wang, S., Robinet, P., Smith, J. D., & Gulshan, K. (2015). ORMDL orosomucoid-like proteins are degraded by free-cholesterol-loading-induced autophagy. Proceedings of the National Academy of Sciences of the United States of America, 112(12), 3728–3733. 10.1073/pnas.1422455112

65. Ye, Y., Shibata, Y., Kikkert, M., van Voorden, S., Wiertz, E., & Rapoport, T. A. (2005). Recruitment of the p97 ATPase and ubiquitin ligases to the site of retrotranslocation at the endoplasmic reticulum membrane. Proceedings of the National Academy of Sciences of the United States of America, 102(40), 14132–14138. 10.1073/pnas.0505006102

66. You, H., Ge, Y., Zhang, J., Cao, Y., Xing, J., Su, D., Huang, Y., Li, M., Qu, S., Sun, F., & Liang, X. (2017). Derlin-1 promotes ubiquitylation and degradation of the epithelial Na+ channel, ENaC. Journal of Cell Science, 130(6). http://jcs.biologists.org/content/130/6/1027.long

67. Zeng, Q., Yao, C., Zhang, S., Mao, Y., Wang, J., Wang, Z., Sheng, C., & Chen, S. (2025). ORMDL3 restrains type I interferon signaling and anti-tumor immunity by promoting RIG-I degradation. ELife, 13. 10.7554/ELIFE.101973

